# Multiscale Hyperbolic Embedding Reveals Hierarchical Structure in Complex Biological Systems

**DOI:** 10.1101/2025.09.29.679407

**Authors:** Mingchen Yao, Anoop Praturu, Tatyana Sharpee

## Abstract

The rapid expansion of biological and computational datasets demands scalable methods that support both visualization and quantitative interpretation. Hyperbolic embeddings are well suited to represent hierarchical structure, but existing approaches are limited by fixed curvature assumptions or poor scalability to large datasets. We introduce MuH-MDS, a multiscale hyperbolic multidimensional scaling algorithm that employs an *adiabatic optimization strategy*: local positions are iteratively refined while cluster centroids are temporarily fixed. This strategy accelerates computation by 10^3^ and enables scaling to datasets with over 80,000 samples. Applied to diverse benchmarks, including *C. elegans* embryogenesis scRNA-seq data, MuH-MDS uncovers intrinsic hierarchical organization and improves both pseudotime inference and lineage reconstruction relative to UMAP and other standard methods. In contrast to UMAP and t-SNE, which prioritize local neighborhoods at the expense of global coherence and metric fidelity, MuH-MDS preserves both local detail and global hierarchy, providing a metrically faithful framework for multiscale analysis of complex biological systems.

## Introduction

Many modern biological datasets, such as gene regulatory networks and single-cell transcriptomes, exhibit intrinsic hierarchical structure [1]. These hierarchies reflect fundamental biological processes, including developmental lineages and molecular cascades where certain genes or molecules exert predominant influence. Traditional Euclidean embeddings, which treat pairwise distances uniformly, often fail to capture these asymmetries and nested relationships [2, 3]. In particular, they struggle to represent the few principal axes along which selective amplification or differentiation occurs.

Hyperbolic space offers a compelling alternative. Like a branching tree, hyperbolic space expands exponentially with radius, so hierarchical depth can be represented naturally by radial distance from the origin. Meanwhile, angular coordinates encode branching structure, separating distinct lineages or functional pathways at the same hierarchical level. With its exponentially expanding number of states, it is particularly well-suited for datasets with hierarchical organization [4, 5]. In such embeddings, hierarchical depth maps directly onto radial distance. For instance, the hyperbolic embedding of *C. elegans* embryogenesis sequencing data reveals the radial axis as an effective proxy for developmental pseudotime [3]. Compared to Euclidean approaches, hyperbolic embeddings reduce distortion in global structure [6], enhancing interpretability and visualization of complex biological hierarchies [5].

Previous work has demonstrated the advantages of hyperbolic embeddings for uncovering hierarchical structures in both graphs [7, 8, 9, 10, 11, 12, 13, 14] and biological data [5, 3, 6]. However, most methods are restricted to fixed dimensions and curvature, or fail to scale to the size of modern datasets.

Advances in sequencing technologies now produce datasets that are large-scale, high-dimensional, sparse, noisy, and feature-correlated. These challenges highlight the need for specialized embedding methods while also providing abundant benchmarks for testing [20, 21, 22, 23]. In scRNA-seq, correlated gene expression implies an intrinsically low-dimensional structure, motivating dimensionality reduction, clustering, and pseudotime analysis [15, 16, 17, 3, 5, 18, 19]. Yet Euclidean methods such as UMAP [24] and t-SNE [25] often distort global relationships, misplacing biologically important samples or introducing artificial patterns. By contrast, hyperbolic embeddings naturally accommodate the tree-like organization of biological processes, reducing distortion and preserving hierarchical relationships.

To address these challenges, we developed **MuH-MDS**, a multiscale hyperbolic multidimensional scaling method. Building on Bayesian hyperbolic multidimensional scaling (BHMDS) [12], MuH-MDS introduces a novel global–local embedding framework that efficiently approximates large distance matrices, improving both scalability and accuracy. This design enables the analysis of bioinformatics datasets at unprecedented scale, facilitating the discovery of biologically meaningful hierarchies. As a metric embedding method, MuH-MDS explicitly preserves pairwise distances, providing more interpretable embeddings. By operating in exponentially expanding hyperbolic space, MuH-MDS is able to preserve complex metric relationships even in very low dimensions. As a result, it can faithfully encode hierarchical data structures in embeddings as low as three dimensions, enabling meaningful visualization without sacrificing metric consistency. This makes MuH-MDS particularly well suited for bioinformatics datasets and for uncovering biologically meaningful hierarchical organization.

## Results

### Multiscale hyperbolic-MDS algorithm

The major challenge in creating scalable embedding methods for non-Euclidean spaces is that these methods can no longer rely on linear operations. Evaluating nonlinear distances across the whole dataset at each step of optimization becomes computationally prohibitive. Some methods use local linearization of the space [26] to get access to linear algebra methods that have been optimized for speed. But this leads to global distortions. In contrast, we propose here to simplify most of the distance calculations by substituting distances between disparate samples with their distance to centers of other clusters (Fig. 1a). The key concept of the multiscale approach is to partition the samples into clusters before embedding and then perform the embedding in a “global-local” hierarchical sequence (Fig. 1b, Methods). In the global embedding step, one sets cluster centroid positions (Fig. 1b, left). Then, samples in each cluster are embedded locally, based on intra-cluster distancing and their distances to the nearest cluster centroids (Fig. 1b, right). Distance is estimated under the assumption that distances between clusters are much larger than distances within clusters, which is guaranteed by most clustering methods. This structure greatly reduces the embedding matrices’ size in each step and allows the local embedding step to be parallelized, significantly enhancing computational efficiency. We tested MuH-MDS and found that it provides a valid and efficient approximation (Methods), thereby making the algorithm scalable to much larger datasets.

**Figure 1:**
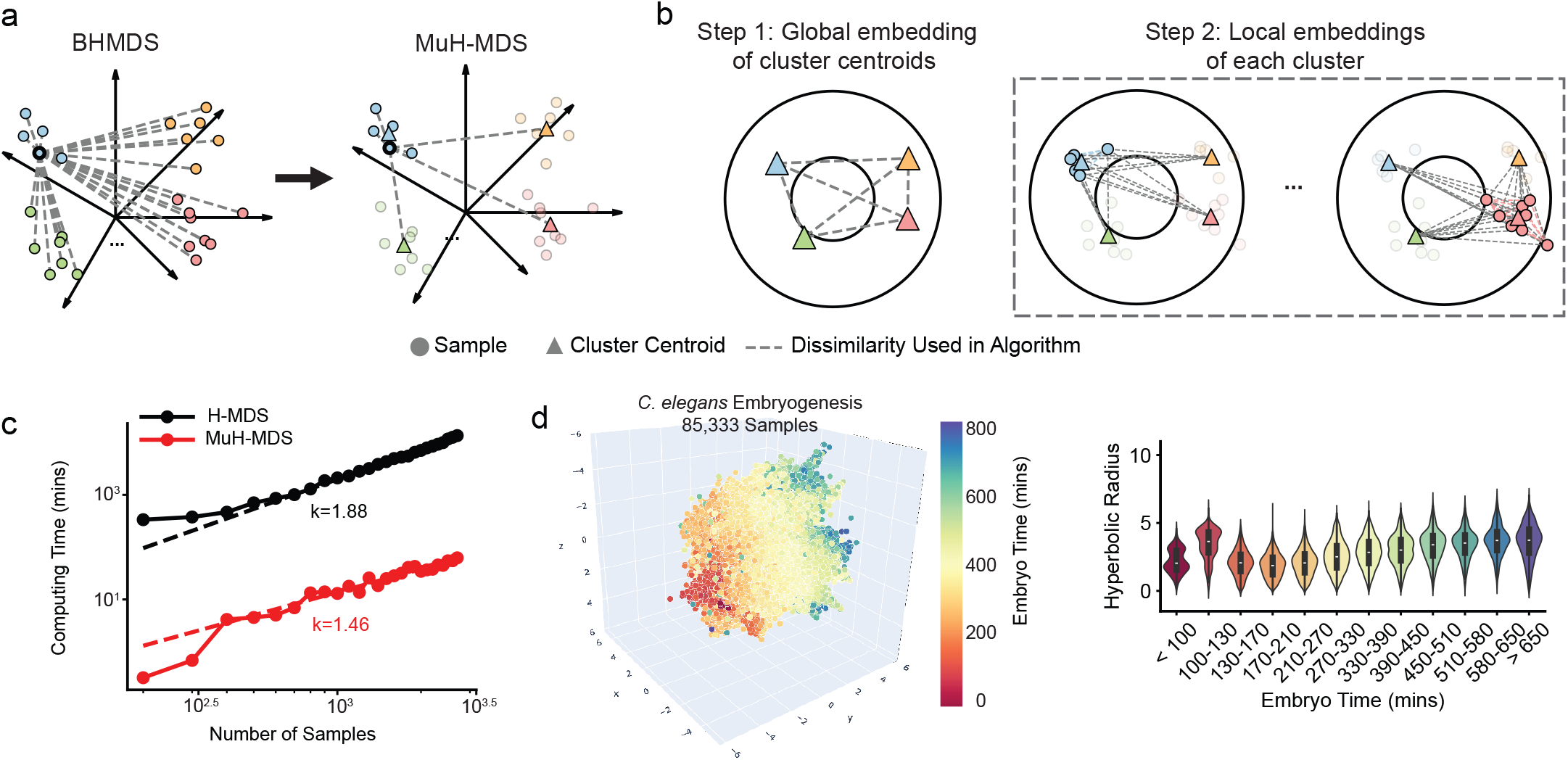
Multiscale hyperbolic multidimensional scaling (MuH-MDS) algorithm. (**a-b**) Schematic representation of (a) the distance approximation and (b) the global-local embedding. (**c**) Comparison of computing time between the original (BHMDS) and multiscale (MuH-MDS) approaches, using the mouse myeloid progenitors scRNA-seq dataset (“Paul”) with 2,730 samples [22]. (**d**) Left: Embedding of the full *C. elegans* dataset (85,333 samples) using the multi-layer approach in 3D hyperbolic space. MuH-MDS parameters: *N*_*neighbors*_ = 30, *N*_*cluster*_ = 800. Right: The embedding preserves the correlation between hyperbolic radius and embryo time in *C. elegans*.

### Scalability and reliability

We compared the performance of MuH-MDS with the previous BHMDS algorithm, whose runtime scales as *O*(*n*^1.88^) with the number of samples *n* (Fig.1c). This behavior is consistent with the expectation given that the number of pairwise distances scales as *O*(*n*^2^). By contrast, MuH-MDS substantially improves scalability, with runtime scaling as *O*(*n*^1.46^) (Fig. 1c), close to the theoretical optimum of 1.33 (Methods).

We assessed embedding quality using metrics such as *Q*_local_, *Q*_global_ [31, 3] (Methods), as well as the correlation between original pairwise distances and embedded distances. For each point, we compute the ranking of all other points according to both the original space and the embedding. At a given neighborhood scale, the *Q* metric measures how consistently points that are close according to the original ranking are also close according to the embedding’s ranking. *Q*_local_ aggregates this agreement over small neighborhood sizes, capturing preservation of local structure, whereas *Q*_global_ aggregates it over larger neighborhood sizes, quantifying preservation of long-range relationships. In contrast, the distance–distance correlation evaluates overall geometric consistency using the raw pairwise distances rather than their rankings, and therefore provides complementary information to *Q*_local_ and *Q*_global_.

Overall, MuH-MDS retained high embedding quality (Fig. S1). While the computing time was reduced by a factor of 10^3^ (from 13,806 minutes for BHMDS to 15 minutes for MuH-MDS), embedding quality metrics only dropped by less than 10% across metrics (Fig. S1c).

MuH-MDS enables the embedding of the full *C. elegans* dataset (85,333 samples) in 1.1 hours on a single CPU. For comparison, the Poincaré map requires 2-3 hours on 1 GPU for 40,000 samples [3]. MuH-MDS perfectly captures the correlation between embryo time and hyperbolic radius (Fig. 1d).

### Robustness to hyperparameter settings

We then evaluated how hyperparameter settings affect the embedding quality metrics and computing time. Several parameters were tested: (1) the number of clusters in the clustering step (Fig. 2a,c), (2) the dimensionality of the hyperbolic space into which the data is embedded (Fig. 2b), (3) the maximum and minimum (Fig. S2a-b) allowed cluster size and (4) the number of nearest neighbors in the local embedding step (Fig. S2c). We found that embedding quality was only sensitive to the number of nearest neighbors for high embedding dimension (Fig. S2c), since more neighboring points are needed as constraints to determine locations in higher dimensionality. Most embeddings intended for visualization purposes are in 2D or 3D spaces, where sensitivity to the number of nearest neighbors can be safely neglected.

**Figure 2:**
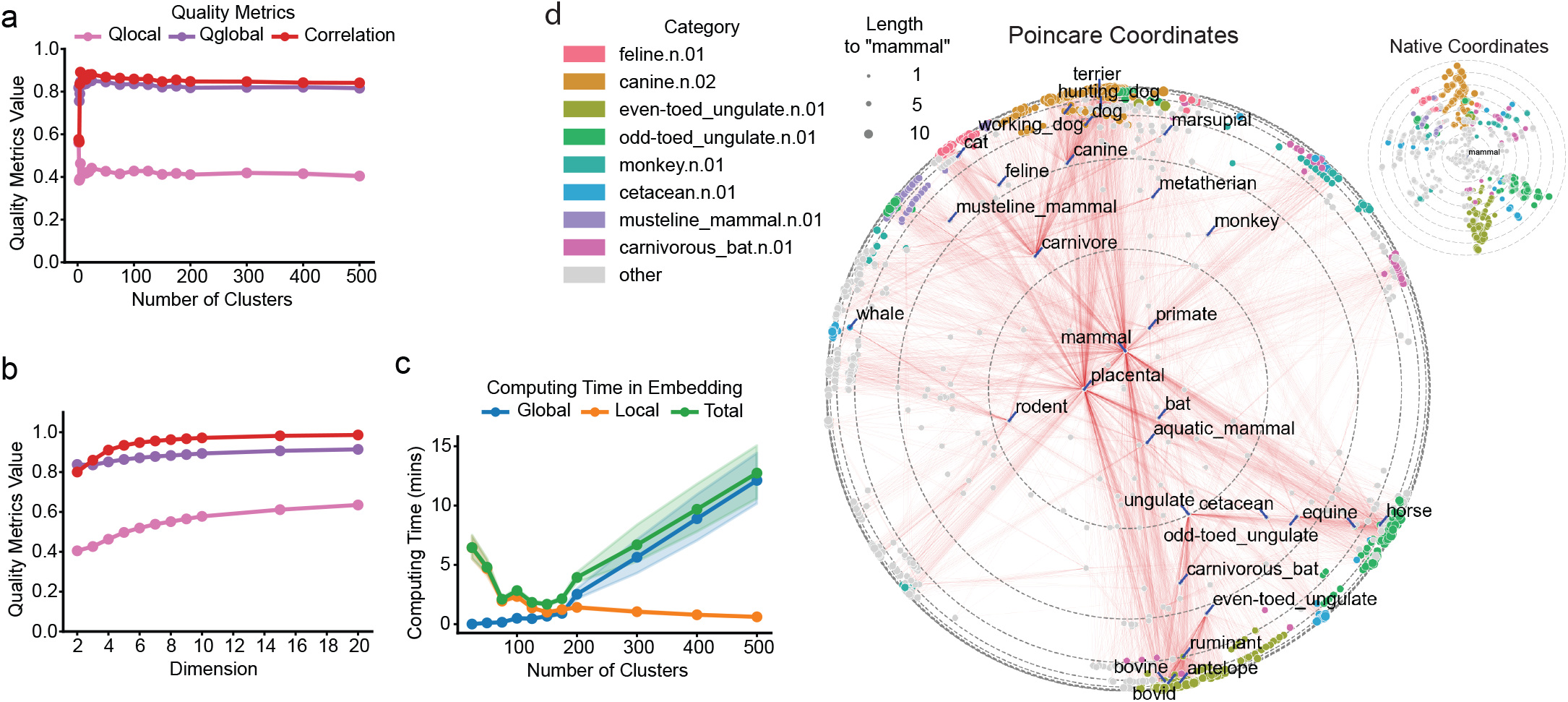
Robustness, dimensional scaling, and hierarchical embedding with MuH-MDS. (**a**) Embedding quality metrics remain stable across different numbers of clusters, except when the cluster number is very small. (**b**) Embedding quality improves with increasing dimensionality of the hyperbolic space, shown here with *k* = 100 clusters. (**c**) Trade-off between global and local computing time for the simulations in (a), with embedding dimension fixed at *D* = 3 for both (a) and (c). All simulations in (a–c) were run with 20 repetitions and *α* = 20 neighbors. (**d**) Application of MuH-MDS to the WordNet mammal subtree. Marker size reflects graph distance from the root node “mammal”, while color indicates membership in semantic categories defined by parent nodes at distance 3 from the root with the largest numbers of child nodes, highlighting major structural branches in the hierarchy. Red lines connect each node to its children. Inset: same embedding in native coordinates, emphasizing child nodes. MuH-MDS parameters: *k* = 198, *α* = 60.

We tested number of clusters (*k*) and found that there is a computing time trade-off between global and local embedding. Embedding quality metrics were sensitive to the number of clusters only when the number of clusters was very small (*k <* 10, Fig. 2a). Beyond this number, embedding quality remained stable, varying by approximately ±5%. This is because *k* ≈ 10 cluster centroids is usually sufficient to determine the global shape of the manifold when embedding into two or three dimensions. By comparison, computing time varied significantly with the number of clusters. There was an optimal number of clusters that minimized computing time. The optimal number of clusters scaled with the number of samples as *k* = *n*^2/3^ to balance the global and local embedding steps. With the optimal number of clusters, the algorithm achieved the best computing time *O*(*n*^1.33^) (Methods). This is consistent with Fig. 1c, where computing time scales as *O*(*n*^1.46^), and with Fig. 2c, where the best computing time is observed between *k* = 100 and *k* = 200, aligning with the theoretical *k* = 2730^2/3^ = 185. Choosing the optimal number of clusters wisely, which can be easily controlled in the clustering step for most clustering methods, significantly optimizes the computing time of MuH-MDS.

We then tested dimension of the space and found that 3D is sufficient to characterize global features. Embedding quality increases monotonically with the dimensionality of the hyperbolic space (Fig. 2b), as higher dimensions provide more capacity to capture details in the data. Notably, while the local embedding quality *Q*_local_ continues to improve with increasing dimensions, the global embedding quality *Q*_global_ and the Shepard diagram correlation saturate at *D* = 4. This indicates that hyperbolic space effectively captures most of the global information within the data in just 2 or 3 dimensions. These results highlight the advantage of hyperbolic embedding for visualizing complex, high-dimensional data in low-dimensional spaces.

To ensure more efficient embedding, we control the size distribution of clusters during the embedding process. For very small clusters, for example, those with only 1-2 samples in a cluster, we apply an “exclude and remap” procedure to reduce the total number of clusters in the global embedding step. Specifically, small clusters are excluded from the global-local embedding process and are mapped to the embedding space only after all other points have been embedded (Methods). The motivation for the “exclude and remap” strategy is that very small clusters provide poor constraints and can degrade centroid-based global structure. By excluding clusters that are too small, the algorithm focuses on capturing the geometry of the majority of points during the embedding step, leading to a more precise representation of the global geometry. For very large clusters that could significantly prolong local embedding time, we further subdivided them into smaller clusters until each sub-cluster size was within the maximum allowed limit. This cluster size distribution control significantly reduced embedding time (Fig. S2b) and helped improve embedding quality (Fig. S2a).

Based on our analyses, we recommend the following hyperparameter choices: (1) set the number of clusters to *k* ≈ *n*^2/3^, where *n* is the number of samples; (2) use a hyperbolic embedding dimension starting at *D* = 3, and increase the dimension only if *D* = 3 is insufficient; (3) set the maximum allowed cluster size to 300–400 samples and the minimum to 3–5 samples; and (4) choose the number of nearest neighbors in the range of 10–40 samples.

### Embedding of graph datasets

While most of our experiments focus on scRNA-seq data, we include several examples of non-biological graph datasets to demonstrate that MuH-MDS performs well across diverse types of data. The classical problem of word embedding serves as an illustrative example of complex network data, where distances are defined as paths in the graph. Using the mammal tree from the WordNet dataset [27], we computed pairwise distances between samples in the graph and performed 2D hyperbolic embedding using MuH-MDS (Fig. 2d). To facilitate comparison with prior studies, we quantify embedding quality using the *distortion rate*. Intuitively, the distortion rate measures how much pairwise distances change after embedding, relative to the original distances. Lower distortion therefore indicates that the embedding more faithfully preserves the overall geometry of the original space. The distortion rate between original graph and our hyperbolic embedding was 0.158, which is comparable to previous embedding methods [10]. Nodes with a smaller distance to the root “mammal” are placed at smaller radius, and vice versa. When represented in native coordinates (Fig. 2d, inset), it is clearer that MuH-MDS gives an embedding that represents node depth in the mammal closure as radial direction, and different categories as angular direction, giving a clear representation of the tree. Our embedding better preserves hierarchical data structures. For instance, feline and canine nodes are positioned close to carnivore, a relationship that was not captured by methods such as HoroPCA [10, 11]. This advantage arises from the metric embedding approach, which maintains the relative scale of distances between samples. Additional results for graph dataset embeddings are shown in Fig. S3. Embedding graph data into a 2D space using MuH-MDS effectively represents the tree-like structure of the graph while preserving original graph relationships with minimal distortion. Compared with HoroPCA [10] (Fig. S3), MuH-MDS obtained comparable or better embedding quality on each dataset.

### Performance evaluation on benchmark bioinformatics datasets

We evaluated MuH-MDS on multiple benchmark scRNA-seq datasets against popular dimensional reduction methods in both 2D and 3D spaces. Results are presented for two classic datasets: the Mouse myeloid progenitors dataset (“Paul”) [22] and *C. elegans* embryogenesis [23] (Fig. 3a-d). In addition, we tested four other datasets with varying hierarchical structures and dataset sizes (Methods; Fig. S4, S7-S21). In both 2D and 3D spaces, MuH-MDS achieves the best global embedding quality and demonstrates high correlation between original distances and embedded distances. In 3D space, MuH-MDS further improves embedding quality both locally and globally, outperforming competing methods. This is consistent with Fig. 2b, where MuH-MDS, as a metric embedding method, shows a substantial performance improvement when increasing the embedding dimensionality from 2D to 3D.

**Figure 3:**
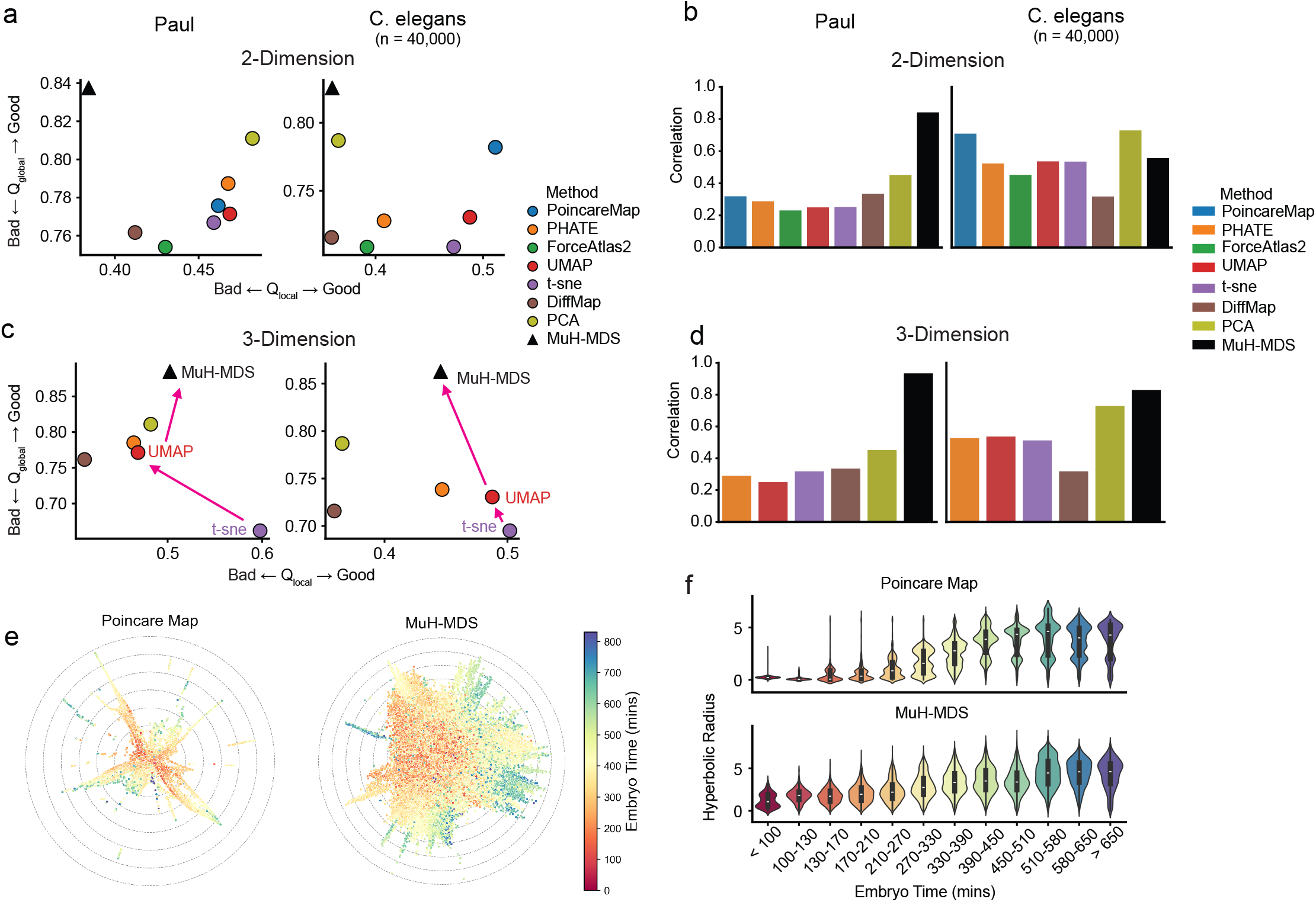
Comparison to existing methods. (**a**-**d**) Embedding quality metrics for two datasets: Paul [22] and *C. elegans* [23], compared with competing methods, in 2D (**a**-**b**) and 3D (**c**-**d**) spaces. (**e**) 2D embeddings of the *C. elegans* dataset (40,000-sample subset) using the Poincaré map and MuH-MDS. (**f**) MuH-MDS more uniformly captures the correlation between hyperbolic radius and embryo time. (See Fig. S4)

To demonstrate the ability of MuH-MDS to uncover hierarchical structures in large-scale data, we analyzed the *C. elegans* embryogenesis dataset, which contains 85,333 samples in total. To ensure scalability for comparison with Poincaré map [3], we randomly sampled 40,000 cells for the analysis in Fig. 3. In this dataset, embryo time is recorded, serving as an ideal indicator of the hierarchy in the differentiation process, where smaller embryo ages correspond to the root of the tree and larger ages to the leaves. The Poincaré map demonstrates that the hyperbolic radius in the embedding correlates with embryo time, reflecting the tree-like structure inherent in the data. MuH-MDS provides a more continuous and well-distributed representation than the Poincaré map (Fig. 3e-f), with lower global distortion (Fig. 3a).

### MuH-MDS reveals hierarchy of cell lineages in *C. elegans* datasets

In bioinformatics, scRNA-seq data offers abundant information about sub-clusters and lineages. However, global structure is often distorted during dimensional reduction with methods such as UMAP when applied to the full dataset [5]. MuH-MDS leverages the properties of hyperbolic space, with its exponential expansion, to effectively capture both global and local structures in the same embedding compared to competing methods.

We tested MuH-MDS on a neuronal progenitor subset, the ABpxp lineage (7,562 samples) from the *C. elegans* dataset. Across varying embedding dimensions, MuH-MDS exhibited higher global embedding quality with dimension 4 and achieved higher local embedding quality starting with dimension 5 (Fig. 4c).

**Figure 4:**
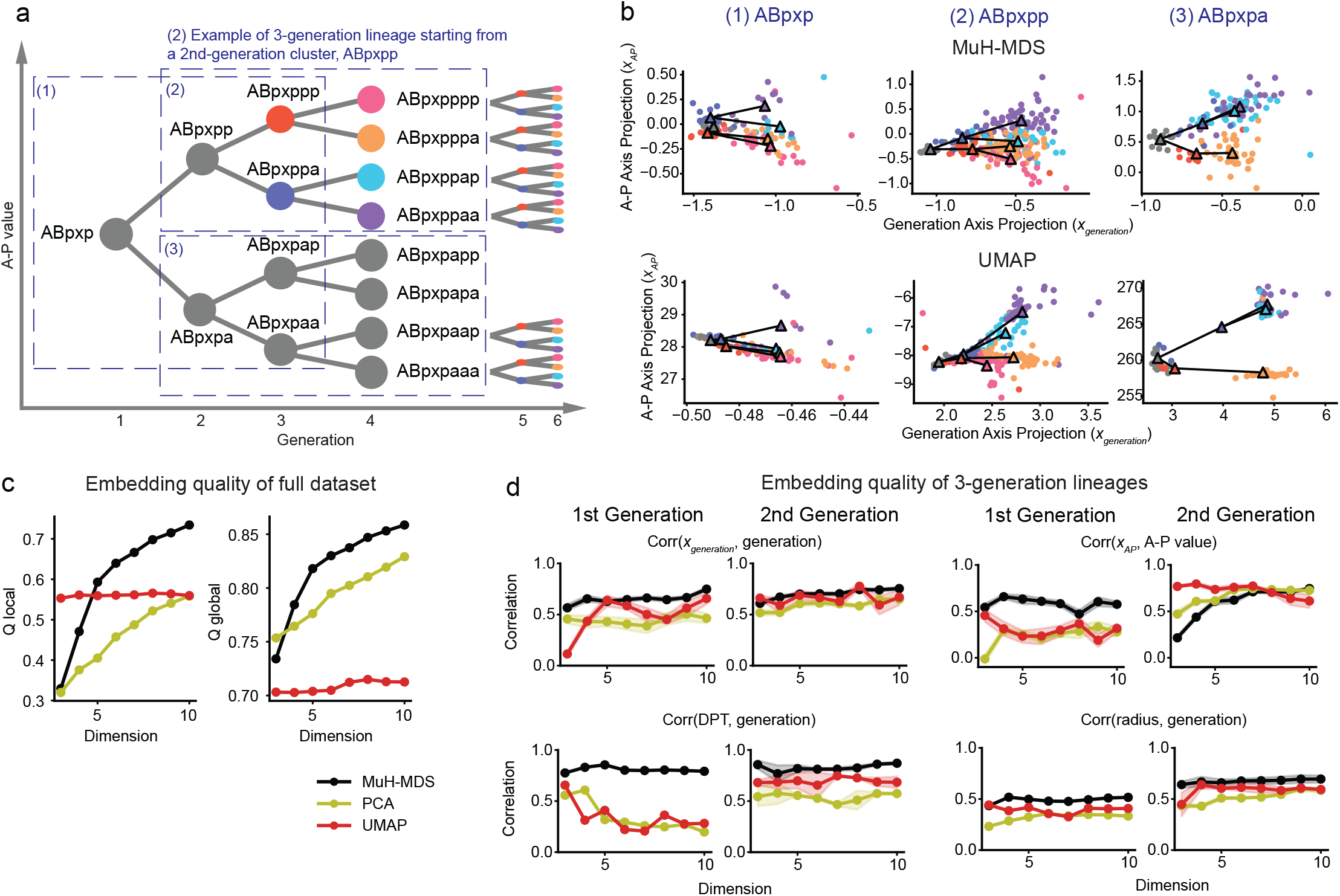
MuH-MDS reveals hierarchical organization in C. elegans ABpxp lineage. (**a**) Schematic of the ABpxp lineage, with anterior (“a”) and posterior (“p”) divisions; dashed boxes mark three sublineages. (**b**) 2D projections from 7D embeddings show MuH-MDS preserves tree-like structure, whereas UMAP merges clusters, such as ABpxppa and ABpxppp (middle column, blue and red triangles) (*α* = 20, k = 50). Triangles indicate cluster centroids of each population, and lines connect the centroids. (**c**) Embedding quality metrics for the full ABpxp lineage (7,562 cells) across dimensions, comparing MuH-MDS, UMAP, and PCA. (**d**) Sub-lineage metrics (generation, “anterior-posterior” correlation (A–P value), pseudotime, and embedding radius) show MuH-MDS consistently outperforms other methods, with improved A–P correlations above dimension 8. See Fig. S5 for more lineages.

To evaluate the preservation of lineage relationships, both locally and globally, we examined 3-generation lineages originating from a single parent node. These parent nodes included global nodes (e.g., ABpxp) and local nodes (e.g., ABpxpppp). Samples within each 3-generation lineage were color-coded based on their relationship to the parent node (Fig. 4a). The representation was visualized in 2D space, mimicking typical visualization procedures, with the embedding projected onto a 1D subspace that best captured either generation depth or differentiation along the “anterior-posterior” axis (“A-P value”) (Fig. 4a, Methods). Samples were randomly split 20 times into 50% training and 50% test sets. For each split, the training set was used to determine the projection direction that best correlated with each metric, and the projection of the test set onto this direction was used to compute the correlation for evaluation. Intuitively, a representation exhibiting high correlations along both projection axes would closely resemble the underlying hypothetical tree structure shown in Fig. 4a, which is exactly what we see in MuH-MDS (Fig. 4b). Meanwhile, UMAP demonstrates strong clustering capabilities but tends to merge distinct and biologically important nodes, such as the ABpxppp and ABpxppa lineages (Fig.4b, middle column).

We computed the correlation between projections onto the generation axis (*x*_generation_) and actual generation depth, and the correlation projections onto the “anterior-posterior” axis (*x*_A-P_) and actual “A-P” value, across different embedding dimensions, for the ABpxp, ABpxpp, and ABpxpa lineages (Fig. 4d; See Fig. S5d for more lineages). MuH-MDS best preserved both generation and “A-P” value across all dimensions for 3-generation lineage from ABpxp, which is the most global lineage (Fig. 4d). It also performed best for ABpxpp and ABpxpa lineages for the generation representation and “A-P” value representation beyond dimension 6. While UMAP performed well for local parent nodes (Fig. S5d, last column), it failed to accurately represent global nodes (Fig. 4d). While methods like UMAP excel in local embeddings, they often distort global structure when applied to the entire dataset. Additionally, UMAP’s non-metric approach results in representations where generation and anterior-posterior relations may not align in biologically meaningful ways. In contrast, MuH-MDS efficiently preserved both global and local hierarchical structures. To better align with standard bioinformatics analyses, we computed pseudotime from each embedding, computed using diffusion pseudotime (DPT) based on the diffusion map constructed from the local neighborhood graph (Methods). DPT derived from MuH-MDS showed the strongest correlation with the ground truth generation depth across all dimensions and lineages. Since the hyperbolic radius can also serve as an indicator of developmental pseudotime, we further computed the correlation between sample radius and generation depth in each embedding. MuH-MDS again exhibited the highest correlation. For more localized lineages, the hyperbolic radius could be a better indicator of developmental progression than DPT (Fig.S5d). This may be because DPT infers temporal order by modeling diffusion processes over a k-nearest neighbor graph constructed from the data. The structure of this graph is sensitive to local neighborhood proximity and the choice of the number of neighbors: too few may lead to disconnected components, while too many may obscure local topology and distort temporal inference. In contrast, the radius offers a more stable geometric measure, as it does not rely on a discrete graph and instead reflects global geometric structure and captures continuous distance from the origin in hyperbolic space. In summary, MuH-MDS is a powerful method for preserving both global and local structures in datasets with complex hierarchical relationships.

### MuH-MDS reveals new lineage in mouse myeloid progenitors dataset

We evaluated MuH-MDS on the mouse myeloid progenitor dataset (“Paul”) [22], which provides an example of a dataset with more ambiguous hierarchical structure than *C. elegans*. This classic single-cell RNA-sequencing dataset profiles mouse hematopoietic progenitor cells and is widely used to study differentiation trajectories and to benchmark pseudotime inference and lineage reconstruction methods. In the original study, the authors identified 19 clusters.

Across embedding dimensions ranging from 2 to 10, MuH-MDS achieved higher global embedding quality at all dimensions and higher local embedding quality starting from dimension 4 (Fig. 5b).

**Figure 5:**
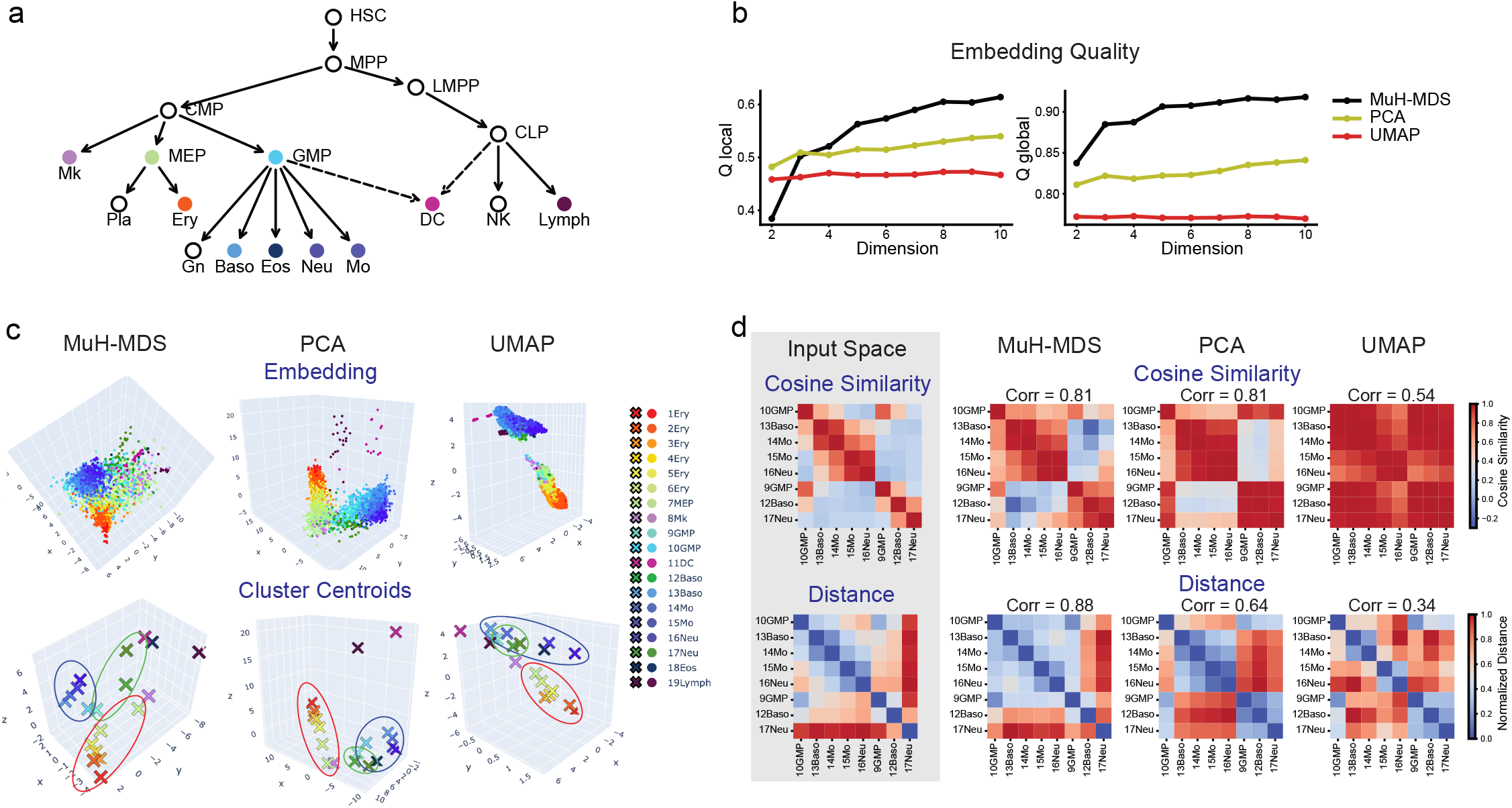
MuH-MDS reveals sub-population in mouse myeloid progenitors dataset. (**a**) Canonical hematopoietic lineage tree [3], where colored nodes represent cell populations defined in the dataset and white nodes denote annotated intermediate states. See Methods for explanations of identified cell type names. (**b**) Embedding quality metrics for the “Paul” dataset across varying embedding dimensions, comparing MuH-MDS, UMAP, and PCA. (**c**) Three-dimensional embeddings produced by MuH-MDS, PCA, and UMAP. Points are colored according to the cluster annotations in the original study [22], and cluster centroids are computed in each embedding space. The three sub-populations discussed in the text are highlighted by colored outlines around the centroids. (**d**) Geometric comparison between the input space (100-PC PCA space derived from the original RNA-seq data) and each three-dimensional embedding. MuH-MDS best preserves both cosine similarity and pairwise distances from the input space. Spearman correlation between the similarity/distance matrices of the embedding space versus the input space is computed. See Fig. S6 for results across all 19 clusters.

PCA and UMAP reveal two major lineages: MEP → Ery (Fig. 5c, middle and right columns; clusters 1–7, red outline) and GMP → {Baso, Mo, Neu, Eos} (Fig. 5c, middle and right columns; clusters 9–18, blue outline). In contrast, MuH-MDS separates three distinct sub-populations, distinguished by angular direction in the embedding space (Fig. 5c, left column): (1) MEP → Ery (clusters 1–7, red outline), (2) 10GMP → {13Baso, 14Mo, 15Mo, 16Neu} (blue outline), and (3) 9GMP → {12Baso, 17Neu} (green outline). In PCA and UMAP, the green sub-population (9GMP, 12Baso, 17Neu) are intermingled with the blue sub-population (10GMP, 13Baso, 16Neu, and other clusters), whereas MuH-MDS clearly separates it along a distinct angular direction in the embedding space. This suggests that these clusters may be more appropriately annotated as a separate sub-population within the hierarchy.

To assess whether this subtle lineage separation is supported by the original data, we compared the geometry of the reduced-dimensional embeddings with that of the input space (100-PC PCA space derived from the original RNA-seq data) (Fig. 5d). Both cosine similarity and pairwise distance analyses show that the 10GMP and 9GMP lineages are already separated in the input space, in terms of both angular relationships and raw distances. Among the three reduced-dimensional representations, 3-dimensional MuH-MDS most accurately captures these subtle sub-lineage differences and best preserves the global pairwise geometry among all clusters (Fig. S6).

Overall, MuH-MDS effectively preserves both global and local structure in datasets with complex hierarchical organization, enabling the resolution of lower-level sub-lineages and providing additional biological insight.

We note that the Poincaré map [3] also recovers this sub-lineage, further illustrating the advantage of hyperbolic embeddings over Euclidean methods in revealing fine-grained local structure. Importantly, MuH-MDS achieves comparable representational power with substantially lower computational cost.

### Biological insights enabled by hyperbolic embeddings

In summary, by metrically embedding data in hyperbolic space, MuH-MDS enables biological insights that are difficult to obtain from Euclidean representations.

In the *C. elegans* embryogenesis dataset, Euclidean embeddings preserve major developmental trends but frequently merge biologically distinct lineages at shallower hierarchical levels. For example, the “ABpxppa” and “ABpxppp” lineages are merged in UMAP representation, but are separated in MuH-MDS (Fig. 4b, left and middle column; blue and red triangles). By operating in hyperbolic space, MuH-MDS encodes developmental depth along the radial direction and branching structure in angular coordinates, resulting in embeddings with clear biological interpretability: both diffusion pseudotime and hyperbolic radius derived from MuH-MDS serve as robust indicators of developmental progression.

Similarly, in the mouse myeloid progenitor dataset, PCA and UMAP recover two dominant differentiation sub-populations (Fig. 5c, middle and right column), but fail to resolve finer sub-population structure within GMP-derived populations. In contrast, MuH-MDS identifies additional GMP-to-Neu and GMP-to-Baso sub-populations that are geometrically distinct (Fig. 5c, left column, green outline) and consistent with separations already present in the original high-dimensional data (Fig. 5d). The recovery of this subpopulation suggests that hyperbolic geometry provides a principled framework for revealing biologically meaningful hierarchical structure that is otherwise obscured in Euclidean embeddings.

## Discussion

Leveraging a multiscale global-local embedding approach, we developed MuH-MDS, a hyperbolic MDS embedding method that achieves both efficiency and scalability on large datasets. MuH-MDS uses adiabatic approximation to find local embedding positions within a cluster while keeping inter-cluster positions fixed. The metric property of the embedding ensures that the low-dimensional representation is quantitatively meaningful, while the well-defined hyperbolic geometry makes the method suitable for further analyses, such as identifying feature-correlated axes, subtypes, and lineages, as well as performing computations between vector representations of samples. The concept of global-local approximation can potentially be generalized to other cases where large dissimilarity matrices require approximation under a nonlinear metric, a scenario that often poses scalability challenges.

Unlike traditional methods like PCA, which provide a well-defined vector space but often distort relationships between samples due to limited dimensionality, hyperbolic embedding addresses this limitation. By leveraging its exponentially expanding space along the radius, hyperbolic embedding effectively characterizes complex hierarchical data within constrained dimensional spaces. When hierarchical structure is weak or absent in the data, MuH-MDS naturally adapts by fitting curvature values close to zero[12], causing embeddings to concentrate near the origin where hyperbolic geometry locally approximates Euclidean space, and thus avoiding the introduction of artificial hierarchy. While MuH-MDS handles zero-curvature datasets, one may benefit less from the hierarchical interpretations of MuH-MDS since the data itself lacks hierarchy. In such cases, Euclidean methods suffice.

We observed a trade-off between global and local embedding times, driven by the balance between cluster numbers and sizes. The main impact on computing time that can be achieved without compromising embedding quality is to select appropriate clustering parameters, with the number of clusters scaling as *n*^2/3^. Implementing additional cluster size distribution controls, such as re-dividing overly large clusters, has been shown to further reduce computing time while maintaining good embedding quality (Fig. S2).

The primary ways for future improvements of the algorithm is to allow for multiple rounds of updates in the centroid position. For current datasets, a single round has been sufficient, cf. Fig. 2b, but as the datasets size increases further, it might become necessary to consider multiple rounds of global updates. Another point that deserves careful consideration is the definition and computation of the original distances. When using a metric embedding algorithm, it is crucial to consider whether the chosen metric appropriately captures the desired features and operates at a reasonable scale. In scenarios where defining a meaningful metric is difficult or only rank-based relationships between samples are of interest, non-metric methods may be more appropriate. We note that the adiabatic approximation employ here to improve metric multi-dimensional scaling can be also be used to improve non-metric multidimensional scaling approaches [28].

## Methods

### Hyperbolic geometry

The literature on hyperbolic geometry in complex systems is vast, and we do not intend to cover it in detail here. We cover only the basics necessary for the model and refer the reader elsewhere for more details [4]. Hyperbolic space can be represented using several equivalent coordinate systems. In this work, we make use of three commonly used representations: native (geodesic polar), Poincaré, and Lorentz coordinates. Although these models differ in their parameterizations and distance formulas, they describe the same underlying geometry and are related through smooth, invertible transformations, emphasizing different aspects of hyperbolic geometry. Throughout this work, we adopt the representation most appropriate for the computational or interpretative task at hand.

In the native coordinate system, a point is represented by a radial coordinate *r* ≥ 0 and angular coordinates *θ*, analogous to polar coordinates in Euclidean space. The radial coordinate *r* corresponds directly to the hyperbolic distance from a chosen origin, while *θ* specifies direction along the manifold. The hyperbolic metric takes the form

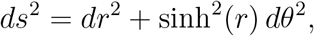

illustrating how distances expand exponentially with *r*. This representation provides an intuitive interpretation of hierarchical structure, where increasing depth in a hierarchy corresponds to increasing radial distance.

The Poincaré coordinate system represents hyperbolic space within a bounded Euclidean domain, such as the unit disk (or unit ball in higher dimensions). Points are constrained to satisfy ‖*x*‖ < 1, and hyperbolic distances are given by

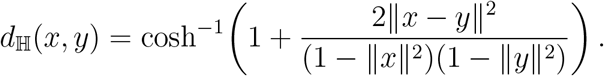

In this model, distances are increasingly distorted near the boundary: Euclidean distances near ‖*x*‖ ≈ 1 correspond to large hyperbolic distances. For visualization, we typically prefer native coordinates because radial distance directly corresponds to hyperbolic distance and hierarchy depth, whereas the Poincaré model compresses large hyperbolic distances near the boundary, making relative depths harder to interpret quantitatively.

The Lorentz coordinate system embeds hyperbolic space into ℝ^*D*+1^ equipped with a Minkowski inner product (See mathematical details in the next paragraph). Although less visually intuitive, the Lorentz model offers numerical stability and simpler expressions for geodesics and gradients, making it well suited for optimization and large-scale embedding.

We follow [2] who found that using Lorentzian coordinates for hyperbolic space is significantly more computationally stable. In this model, a *D*-dimensional hyperbolic space is represented by a future facing space like sheet in a *D* + 1-dimensional Minkowski space-time, i.e., the set of points satisfying

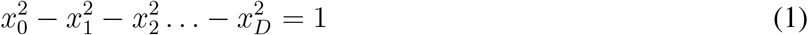

introducing the metric tensor *g*_*αβ*_ = Diag(1, −1, −1,…, −1), where Greek indices run from 0 to *D*, and employing the Einstein summation convention where repeated indices are summed over, we can compactly express the constraint equation as *g*_*αβ*_*x*^*α*^*x*^*β*^ = 1. Continuing with this notation, the distance between two points *x*^*α*^ and *y*^*β*^, which satisfy the constraint equation, is

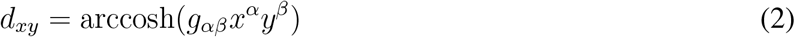

In practice, the *D* space-like components 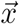 of the coordinates were taken as free parameters, and the time-like component *x*_0_ was computed according to the constraint equation 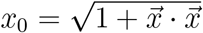.

While coordinates in the model are with unit curvature *K* = −1, the curvature can be effectively fit by fitting for an overall scaling factor of the embedded distance matrix. This is because in curved spaces, the strength of the effect of curvature depends on the scale at which you observe the data. We refer the reader to [12] for the mathematical details of this approach.

### Bayesian hyperbolic multidimensional scaling algorithm

MuH-MDS is based on a Bayesian Hyperbolic MDS algorithm (BHMDS) [12]. Briefly, the generative model assumes the data *δ*_*ij*_ are generated from a geometric distance matrix *d*_*ij*_ in hyperbolic space by a noisy process, and subject to rescaling:

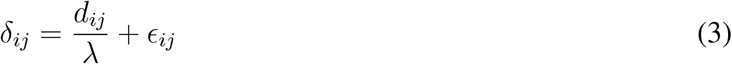

Where *ϵ*_*ij*_ ∼ 𝒩 (0, *σ*_*ij*_) are independent normally distributed variables with mean 0 and possibly differing variances. In practice, *δ*_*ij*_ was normalized so that max(*δ*_*ij*_) = 2, so *λ* corresponds to the radius of the distribution of points in hyperbolic space and probes the effects of curvature (See [12] for more details on this). From this generative model, the likelihood is

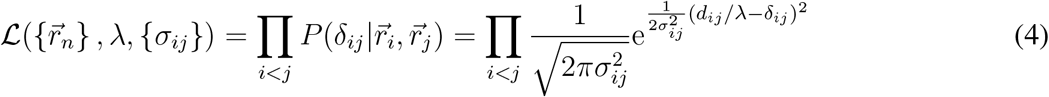

As discussed by BHMDS [12], the number of uncertainty parameters was reduced by assuming each point has a characteristic uncertainty *σ*_*i*_ with an inverse gamma prior, and compute 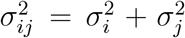. This has an interpretation in terms of springs connected in series [12]. We assume a flat prior for the embedding coordinates and put a normal prior on *λ*. Since the log-likelihood scales as *N*(*N* − 1)/2, the log prior on *λ* was multiplied by *N*(*N* − 1)/2 so that the strength of the prior is not reduced in the large *N* limit. Combining all of this, the negative log-posterior, i.e., loss function, is

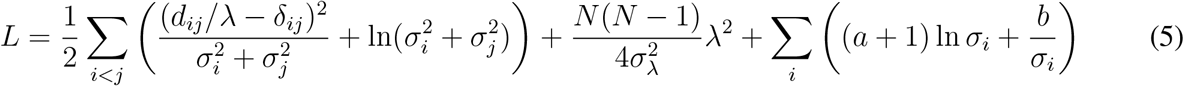

Where in practice [12] we take *σ*_*λ*_ = 10, *a* = 2, and *δ* = 0.5. All parameters were fitted using an LBFGS optimizer implemented in the Stan statistical programming package.

### Global-local embedding procedure in the MuH-MDS algorithm

Step 1: Clustering. Before embedding, samples are divided into different clusters using classical clustering methods, such as k-means clustering. Denote a sample in *L*-dimensional feature space as 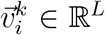, where *i* denotes the sample index and *k* denotes the cluster it is assigned to. For cluster *k*, the cluster centroid was computed as 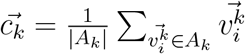, where *A*_*k*_ is the set of samples in cluster *k*.

Step 2: Compute distance matrices. The “global” embedding refers to the embedding of cluster centroids using traditional BHMDS, for which we need the distance matrix between cluster centroids 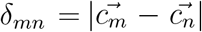. By definition, *δ*_*mn*_ is a *K*-by-*K* symmetric matrix, where *K* is the number of clusters. The “local” embedding fits each individual cluster *A*_*k*_ using previously embedded centroid locations, for which we need two types of distances: (1) the distance between samples in cluster *A*_*k*_, which is computed as 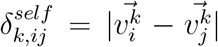 symmetric matrix; (2) the distance from samples in cluster *A*_*m*_ to its nearest *α* neighboring cluster centroids, which is computed as 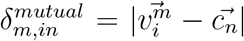. Here, 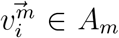, and 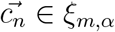 where *ξ*_*m,α*_ is the set of *α* nearest neighbors of 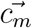. Therefore, 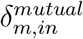 is a |*A*_*m*_|-by-*α* asymmetrical matrix.

The above-mentioned approach provides computations of distance matrices from the feature matrix 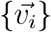. In some cases, such as computing path length in a graph, the feature matrix is unknown, and the relation between samples was characterized directly by the distance matrix. Therefore, we also provide an approach to estimate the *δ*_*mn*_ and 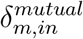 from the original distance matrix (Supplementary References).

Step 3: Global embedding. Global embedding performs BHMDS on *δ*_*mn*_ and gets low-dimensional representations of cluster centroids in *D*-dimensional hyperbolic space ℍ^*D*^. Denote the optimized cluster centroids in ℍ^*D*^ as 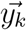.

Step 4: Local embedding. For each cluster *m*, local embedding is performed with a modified version of the BHMDS optimization (the “Relax” optimization): given a set of existing fixed locations 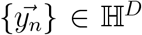, embed a new set of samples optimizing over only their distance matrix 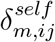 and their distances to existing points 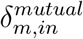. Therefore, instead of summing over all sample pairs (*i, j*) in eq. 5, the summation term in the new objective function is approximated by two terms: ∑_*i,j*_ → (∑_*im,jm*_ + ∑_*im,n*_). The first term is a summation of distances between new samples (in this case, samples in *A*_*m*_), and the second term is a summation of distances between new samples and existing points. It is worth noticing that all the cluster centroid locations needed for local embedding have been computed in the first step, which means local embedding can be parallelized across clusters.

Step 5: Integrate embedding results. From each local embedding of cluster *k*, the optimized locations of individual samples in ℍ^*D*^ were obtained as 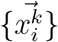. Combining all embedding results, 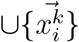 gives the final embedding positions of all samples 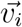 in ℍ^*D*^.

Step 6: Map outliers. When the cluster size and number control procedure is applied, clusters smaller than a specified threshold are excluded from both the global and local embedding steps. Once the embedding results for the remaining data points have been integrated, the previously excluded outliers are remapped into the embedding space using an approach similar to the local embedding step. Given the embeddings 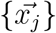, each “outlier” 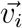 is mapped to the space based on its distance to its nearest neighbors, 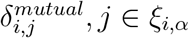. After this step, the embedded outliers are integrated with the previously embedded points, resulting in the final embedding of all samples.

### Proof of the dissimilarity matrix approximation in MuH-MDS algorithm

The global-local embedding procedure provides an effective approximation to the original distance matrix *δ*_*ij*_. Denote the actual distance being optimized in global-local approach as 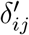. Consider the following three types of sample pairs: (1) sample pair within cluster; (2) sample pair between nearest neighbor clusters; and (3) sample pair between clusters that are not nearest neighbor clusters.

#### Within cluster

By definition of local embedding process, distance within cluster is exactly optimized:

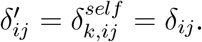

#### Between nearest neighbor clusters

By definition of local embedding process, the distance *δ*_*ij*_ between samples belonging to nearest neighbor clusters was approximated as 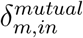, the distance between sample 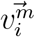 and the cluster centroid 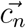 which 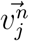 belongs to. Denote 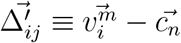, then 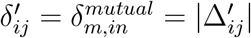.

Given that the samples 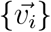 are grouped into clusters 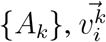 could be written as a displacement from the cluster centroid that it belongs: 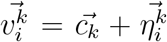. The displacement within cluster is much smaller than the displacement between cluster centroids, giving 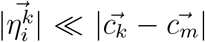, *δm*. Therefore, the true distance *δ*_*ij*_ between a sample 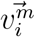 to another sample 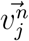 belonging to its nearest neighbor cluster *A*_*n*_ is:

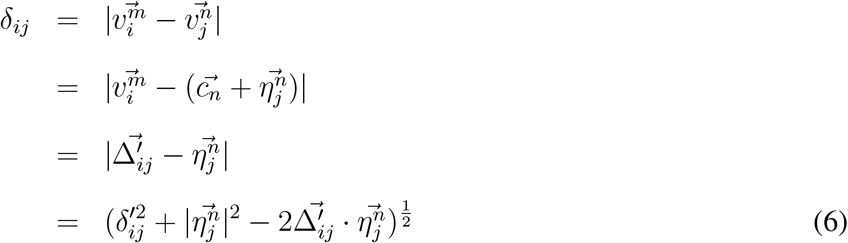

Since 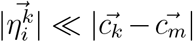, we have 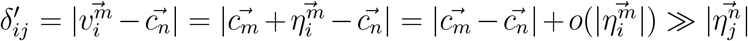. Therefore, 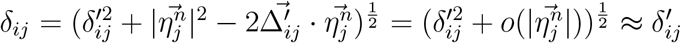.

#### Between clusters that are not nearest neighbor clusters

In this case, the true distance *δ*_*ij*_ could be written as 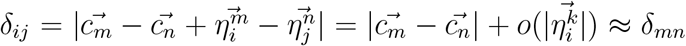, based on similar arguments since that 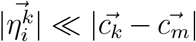. On the other hand, considering the embedding space ℍ^*D*^ and embedded cluster centroids 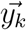 and samples 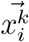, the equivalent distance approximated by global-local process 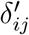 is the distance between 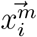 and 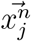. Leveraging the Poincaré disk model [26], we have 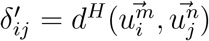, where

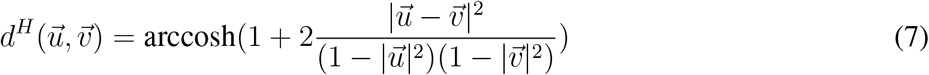

is the distance and 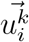 is the representation of 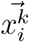 in the Poincaré disk model. Similarly, we denote the representation of 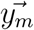 in the Poincaré disk as 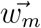. In the global embedding, 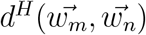 is directly optimized from *δ*_*mn*_, giving 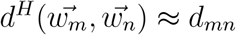. In the local embedding, 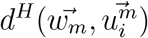 is directly optimized, given that 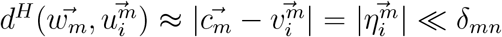. Therefore, 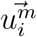 can also be written as a small perturbation to the centroid location: 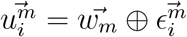, where

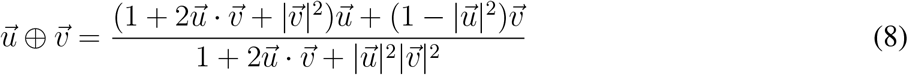

is the Möbius addition, vector addition of points in the Poincaré ball model. Calculating eq. 8, we got 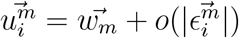. Plugging this into eq. 7 gives: (Supplementary References)

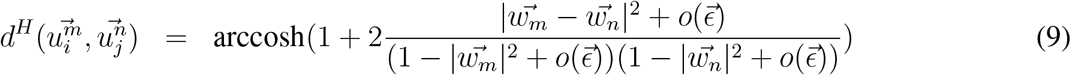

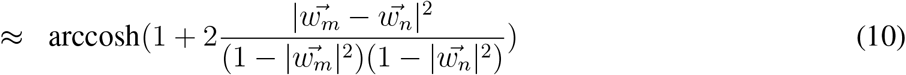

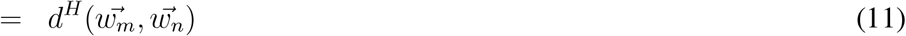

Therefore, 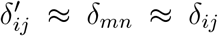. The intuition behind this is that distance between samples belonging to non-nearest-neighbor clusters are not directly optimized. Instead, this distance is approximated by fixing positions of cluster centroids in the global step, and anchoring samples to their nearest clusters in the local step, assuming that local distance is sufficient to constrain the position of samples in low dimensional space.

### Computing time estimation

In the paper, all simulations were conducted on a single x86-64 CPU without parallelization.

Suppose there are *n* samples divided into *k* clusters, the average cluster size is 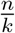. Then the global computing time *t*_*global*_ ∼ *O*(*k*^2^).

#### Without parallelization

When local embedding is not parallelized, local embedding time 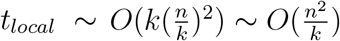.

When 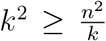, then *k* ≥ *n*^2/3^, computing time is dominated by *t*_*global*_∼ *O*(*k*^2^) with the minimum obtained at *k* = *n*^2*/*3^, which is *O*(*n*^4*/*3^).

When 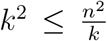, then *k* ≤ *n*^2/3^, computing time is dominated by 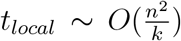 with the minimum obtained at *k* = *n*^2/3^, which is also *O*(*n*^4/3^).

#### With parallelization

When local embedding is parallelized, local embedding time 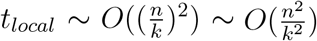.

When 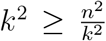, then *k* ≥ *n*^1/2^, computing time is dominated by 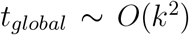 with the minimum obtained at *k* = *n*^1/2^, which is *O*(*n*).

When 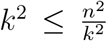, then *k* ≤ *n*^1/2^, computing time is dominated by 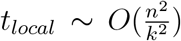 with the minimum obtained at *k* = *n*^1*/*2^, which is also *O*(*n*).

### Dataset descriptions

#### scRNA-seq datasets

For datasets with feature numbers larger than 100, we use PCA to reduce the dimension to 100 before applying the embedding, as well as other competing methods. This is widely used to denoise the data, given the sparsity of single-cell datasets.

#### Toggle Switch

Synthetic data obtained by simulating a simple toggle switch [20] using Scanpy [29], a widely used large-scale single-cell gene expression data analysis tool. It contains 200 samples with 2 markers characterizing a differentiation process with two branches.

#### Myeloid Progenitors (Krumsiek11)

Synthetic data obtained from simulating differentiation of myeloid progenitors, using Scanpy [29]. It contains 640 samples with 11 markers representing cell differentiation of a common myeloid progenitor state to four cell fates: erythrocyte, neutrophil, monocyte, and megakaryocyte.

#### Mouse Myelopoesis (Olsson)

The mouse myelopoesis dataset [21] contains 382 samples belonging to 9 cell types: HSCP-1 (hematopoietic stem cell progenitor), HSCP-2, megakaryocytic, erythrocytic, Multi-Lin (multi-lineage primed), MDP (monocyte-dendritic cell precursor), monocytic, granulocytic and myelocyte (myelocytes and metamyelocytes). We downloaded preprocessed data from Klimovskaia et al. [3], with 382 samples and 533 features.

#### Mouse myeloid progenitors dataset (Paul)

The mouse myeloid progenitors dataset is a MARS-seq dataset from Paul et al. [22]. We loaded from Scanpy and preprocessed using the pipeline offered by Scanpy (sc.pp.recipe zheng17()). After preprocessing, there are 2730 samples and 999 genes kept. The original paper identified 19 clusters.

Abbreviations used in the plots are as follows: HSC, self-renewing hematopoietic stem cell; CMP, common myeloid progenitor; MEP, megakaryocyte/erythrocyte progenitor; GMP, granulocyte/macrophage progenitor; Mk, megakaryocyte; Ery, erythrocyte; Mo, monocyte; DC, dendritic cell; Neu, neutrophil; Baso, basophil; Eos, eosinophil; NK, lymphoid progenitor; and Lymph, lymphocyte.

#### C. elegans embryogenesis

The *C. elegans* dataset is 10X Genomics data from Packer et al. [23]. It contains transcript data of single cells from *C. elegans* embryos at developmental stages ranging from gastrulation to terminal cell differentiation, with 37 main cell types, and embryo time ranging from 0 to 830 minutes. The data is publicly available at [GEO:GSE126954](https://www.ncbi.nlm.nih.gov/geo/query/acc.cgi?acc=GSE126954). The original dataset has 89701 samples and 20222 genes. We preprocessed it using a standard Scanpy [pipeline](https://scanpy-tutorials.readthedocs.io/en/latest/pbmc3k.html) for 10x genomics. After preprocessing, there are 85,333 samples with 2756 genes. Since competing methods like Poincaré map take a very long time to run for large datasets, we randomly sampled 10,000 and 40,000 samples from the full dataset to create smaller datasets for benchmarking. To analyze cell lineages, we create another subset only containing the ABpxp ectodermal lineage, where the descendants are labeled by attaching either “p”(posterior) or “a”(anterior) (ABpxpa, ABpxpp, ABpxpaa, ABpxpap, etc. See Fig. 4a). The ABpxp lineage dataset has 7,562 samples with 2,756 genes, divided into 72 different lineages.

#### WordNet

WordNet [27] is a large lexical database of English. We embedded the mammal subtree of WordNet to demonstrate that MuH-MDS can be generally applied to a variety of datasets. We took the graph data of the mammal subtree from Nickel and Kiela [7] and computed the distance matrix based on the graph path length where each node is only connected to its direct ancestor.

#### Graph Datasets

We evaluated our method on four graph datasets originally introduced by De Sa et al. and obtained data from Chami et al. [10]. These datasets represent diverse hierarchical structures and have been widely used in prior work on hyperbolic embedding. Specifically, they include: a fully balanced tree with 40 nodes, a phylogenetic tree with 344 nodes, a biological disease graph representing disease relationships, with 516 nodes, and a Computer Science Ph.D. advisor–advisee network containing 1025 nodes. We directly obtained the edge list files from the repository and constructed the undirected graph for each dataset. For embedding purposes, we computed the shortest path (graph geodesic) distance matrix between all pairs of nodes in each graph. This distance matrix was then used as input for our embedding algorithm.

### Competing Methods

We compared with competing methods. Details can be found in the code released.

#### Poincaré Map

We compared with Poincaré Map using the code provided by the authors [3] from [GitHub](https://github.com/facebookresearch/PoincareMaps). All datasets were simulated based on the example usage parameters.

#### Phate

We compared with Phate [18] using phate 1.0.11. Using default parameters, we embedded data into either 2 or 3 dimensions. The documentation of Phate can be found here: [link](https://github.com/KrishnaswamyLab/PHATE).

#### Scanpy

Results for the following competing methods: diffusion map [15], UMAP [24], T-sne [25], PCA, ForceAtlas2 [19] were fitted using scanpy (scanpy 1.9.8), a single cell analysis tool [29]. T-sne and ForceAtlas2 are only in 2-dimension, and other methods are in either 2- or 3-dimension. Scanpy Documentation: [link](https://scanpy.readthedocs.io/en/stable/%20).

### Embedding quality metrics

To quantify the embedding performance, we computed a scale independent quality criteria proposed by Lee at al [31]. For each point, the rankings of all other points by distance are computed in both the original space and the embedding. At each neighborhood scale, the metric measures how consistently points that are close according to the original ranking are also close according to the embedding’s ranking. After that, *Q*_local_ summarizes this ranking agreement over small neighborhood scales, reflecting how well local relationships are preserved, whereas *Q*_global_ summarizes it over larger neighborhood scales, quantifying the preservation of long-range structure. Klimovskaia et al. presented detailed calculation of *Q*_*local*_ and *Q*_*global*_ and publicly available [code](https://github.com/facebookresearch/PoincareMaps/blob/main/embedding_quality_score.py).

In WordNet embedding, we computed the distortion rate using the following formula [10]:

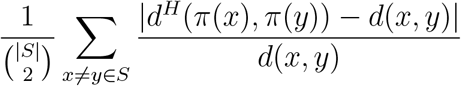

where *d*^*H*^(*a, b*) is the hyperbolic distance between *a, b ∈* ℍ ^*D*^, *π*(*x*) is the hyperbolic embedding of *x*, and *d*(*x, y*) is the original distance.

### Cell lineage data analysis

#### Logarithmic map

Logarithmic map maps a point *x* in the hyperbolic space to a vector in the tangent plane of a given point *p* such that the vector norm of log(*x*) is equal the distance from *p* to *x*. The mapping to tangent space helps generalize matrix and vector calculations into hyperbolic space [26]. For *p* at the origin, so *p*^*α*^ = (1, 0, 0,…, 0), the logarithmic map of *x*^*α*^ is given by

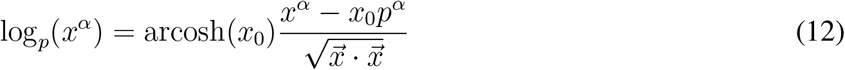

To visualize lineages embedded in 7-dimensional hyperbolic space, we use logarithmic map to project them onto 2-d tangent space. In practice, we used another valid form in Poincaré disk to calculate the vector in tangent space [26]: 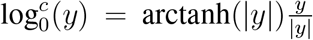. In tangent space, Euclidean vector and matrix computations could be performed.

#### Fitting projection axis

To evaluate how well the learned representation preserved biologically meaningful axes, we performed repeated correlation-based projection analysis. Specifically, for each of 20 repetitions, samples were randomly split into 50% training and 50% test sets. In each split, the training set was used to fit a one-dimensional projection direction that maximized correlation with either generation depth or anterior–posterior (A–P) position, using L-BFGS-B optimization in Euclidean space. The learned projection was then applied to the held-out test set, and the correlation between projected coordinates and the corresponding ground truth labels was computed for evaluation.

#### Diffusion pseudotime (DPT)

DPT was computed using the sc.tl.diffmap and sc.tl.dpt functions in Scanpy [29]. For each lineage, cells were embedded into a diffusion map constructed from a k-nearest neighbor graph (with k=10) based on the input representation. The root cell was selected as the one closest to the centroid of a designated early lineage population, and pseudotime values were then inferred from diffusion components.

## Acknowledgements

This work was funded by the Brown Foundation Research Fellowship to M.Y., and the following grants to T.S.: Paul G. Allen Frontiers Group (19PABH134610000); National Science Foundation (NSF) grants IIS-1724421 and PHY-4213080; the NSF Next Generation Networks for Neuroscience Program (award 2014217); and the Edwin K. Hunter Chair in Computational Neuroscience.

## Author contributions

M.Y., A.P. and T.S. conceived and designed the study. M.Y and A.P. designed the algorithm. M.Y. implemented the method, conducted the experiments, and analyzed the results. M.Y., A.P., and T.S. wrote the manuscript.

## Competing interests

The authors declare no competing financial or non-financial interests.

## Data availability

The pre-processed datasets used in this study are available at https://github.com/ymch815/Multiscale-HMDS. The C. elegans scRNA-seq data is publicly available at GEO:GSE126954.

## Code availability

The source code for MuH-MDS, including the algorithm and scripts for generating all figuresis, is available at https://github.com/ymch815/Multiscale-HMDS.

## Supplementary materials

### Pseudo code for MuH-MDS

Algorithm 1 shows the M-HMDS algorithm. In line 6, *δ*_*ij*_ is a pairwise distance matrix of samples in cluster *m* and cluster *n*; *MInd* and *NInd* indicates which samples belongs to cluster *m* or *n*. In line 13, *d*_*i*,{*n*}_ is a pairwise distance matrix of sample *i* combined with samples in cluster *n*. In line 19, 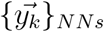 is set of locations of cluster centroids of the nearest neighbors of cluster *m*. In line 25, 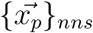 is locations of nearest neighboring points of the outlier 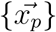, and 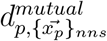 is mutual distance from outliers to them.

#### Algorithm 1

Multiscale hyperbolic multidimensional scaling (MuH-MDS)

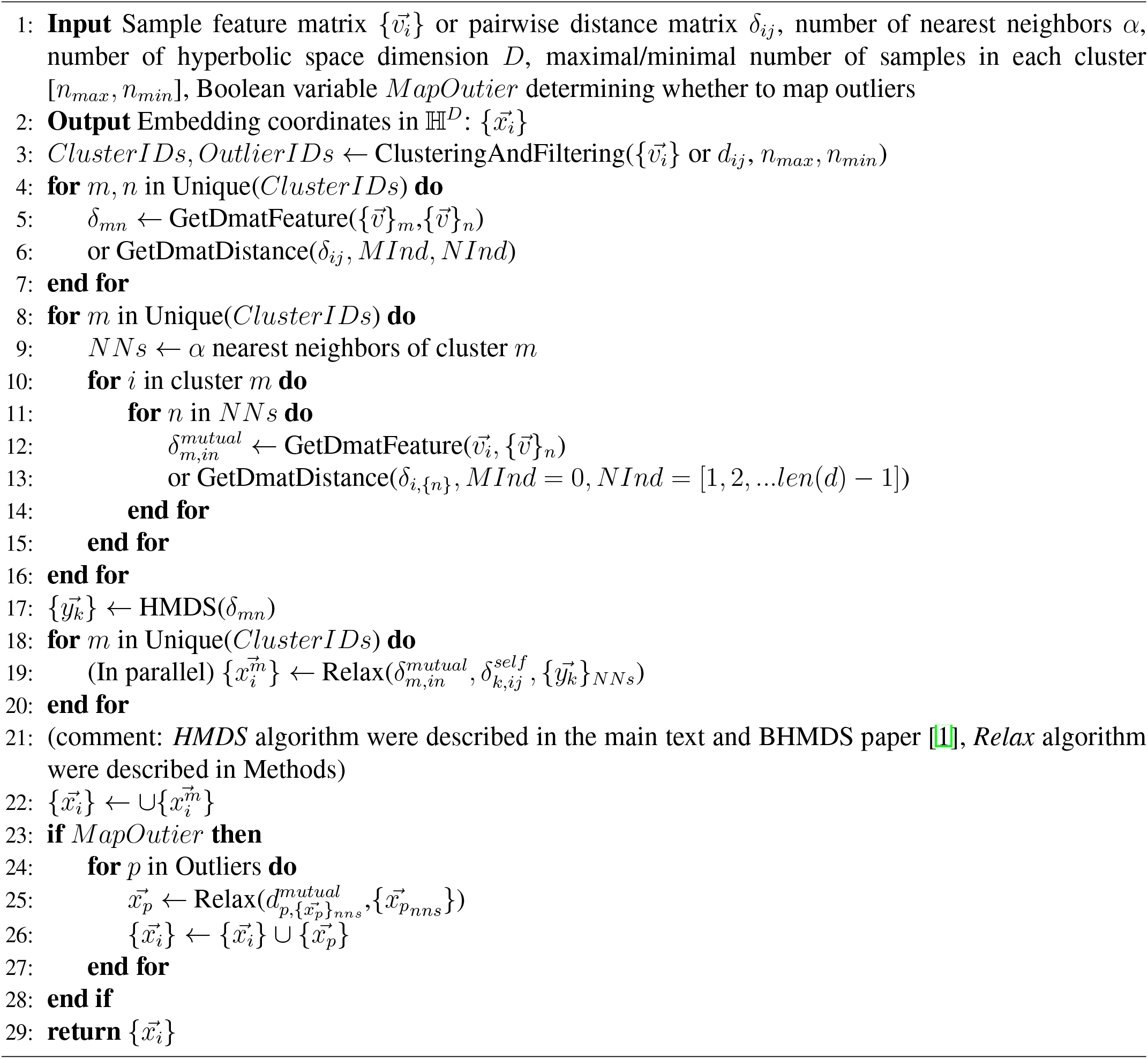

Algorithm 2 simply defines the function for clustering, filtering and cluster size control.

#### Algorithm 2

ClusteringAndFiltering(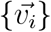 or *δ*_*ij*_, *n*_*max*_, *n*_*min*_)

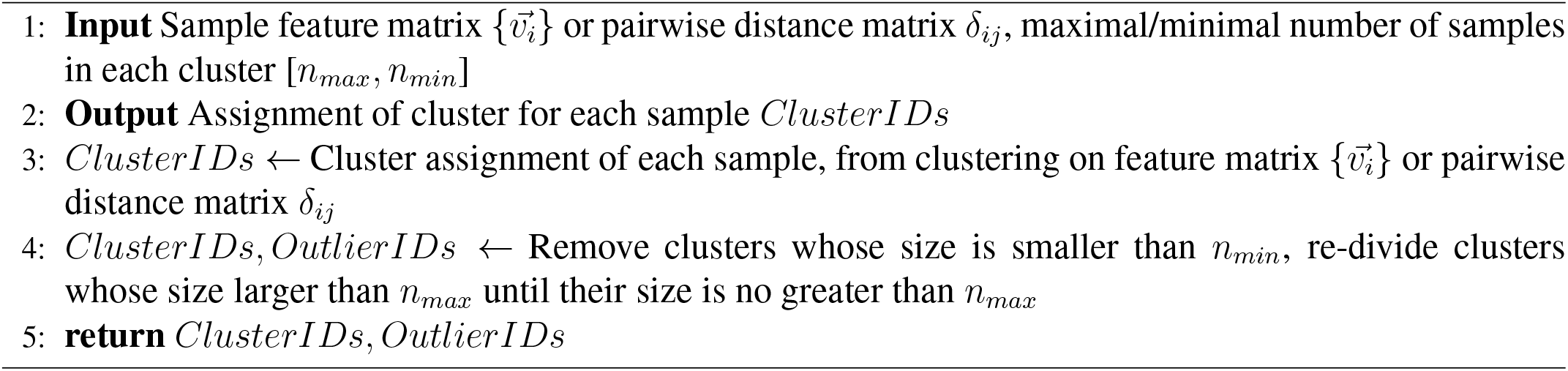

### Compute distance to clusters from distance matrix

Algorithm 3 and 4 defines two different functions to fill the needs for different input types: feature matrix or pair-wise distance matrix. In Algorithm 1, it will determine which distance computing function to implement depend on the input type.

#### Algorithm 3

GetDmatFeature 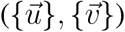

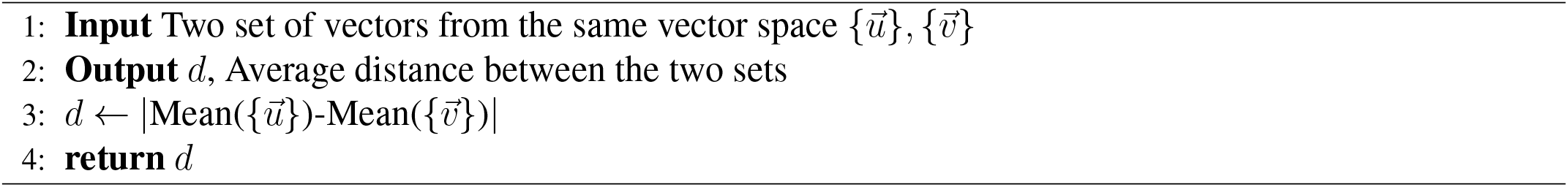

#### Algorithm 4

GetDmatDistance

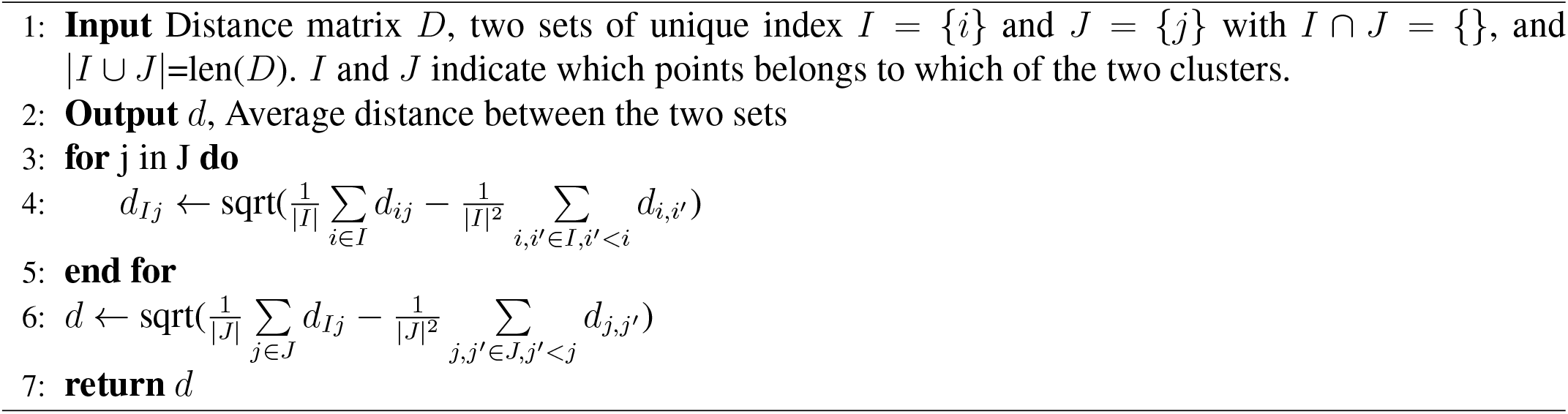

The computation in Algorithm 4 is based on following approximations: Suppose we have a set of *n* vectors 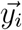 and a vector 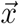. Let 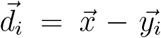. The goal is to approximate 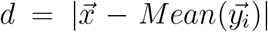 from 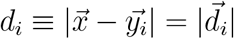 and 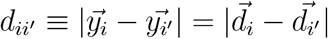.

We have

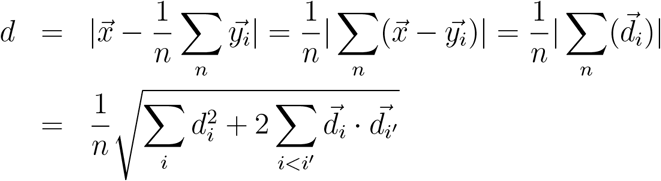

Since 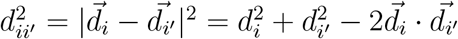, we have 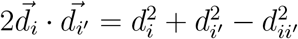. Therefore:

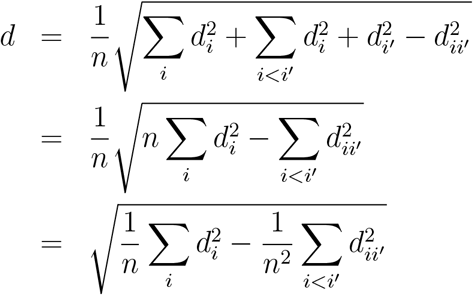

One might notice that Algorithm 4 depends on the order of performing summation on different clusters. We set the order by the variance of original local distance matrix, and sum over cluster with smaller variance first.

### Approximation on distance between clusters that are not nearest neighbor clusters

The Möbius addition is:

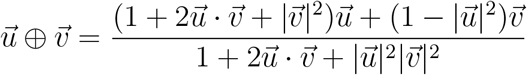

Therefore,

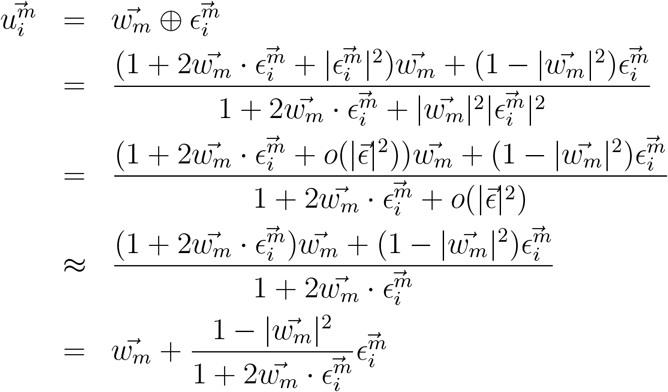

Let 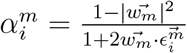, then 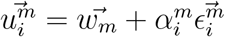, upon dropping the *o*(*ϵ*^2^) term.

Consider the scale of 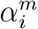: Since 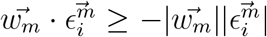, we have 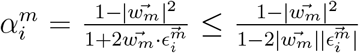. Given that 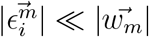, so 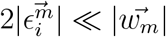, we have 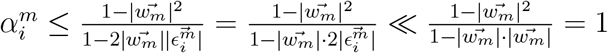.

Since 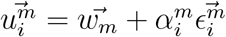 and 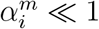, we have 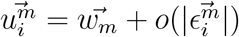.

The distance in Poincaré disk model is:

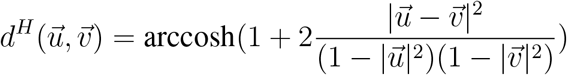

Therefore,

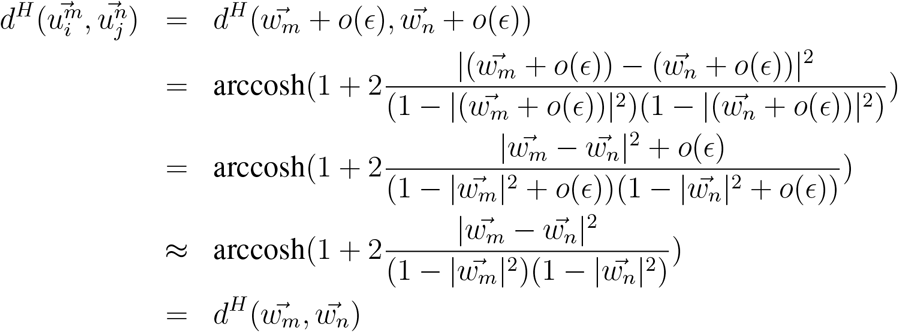

---

**Figure S1:**
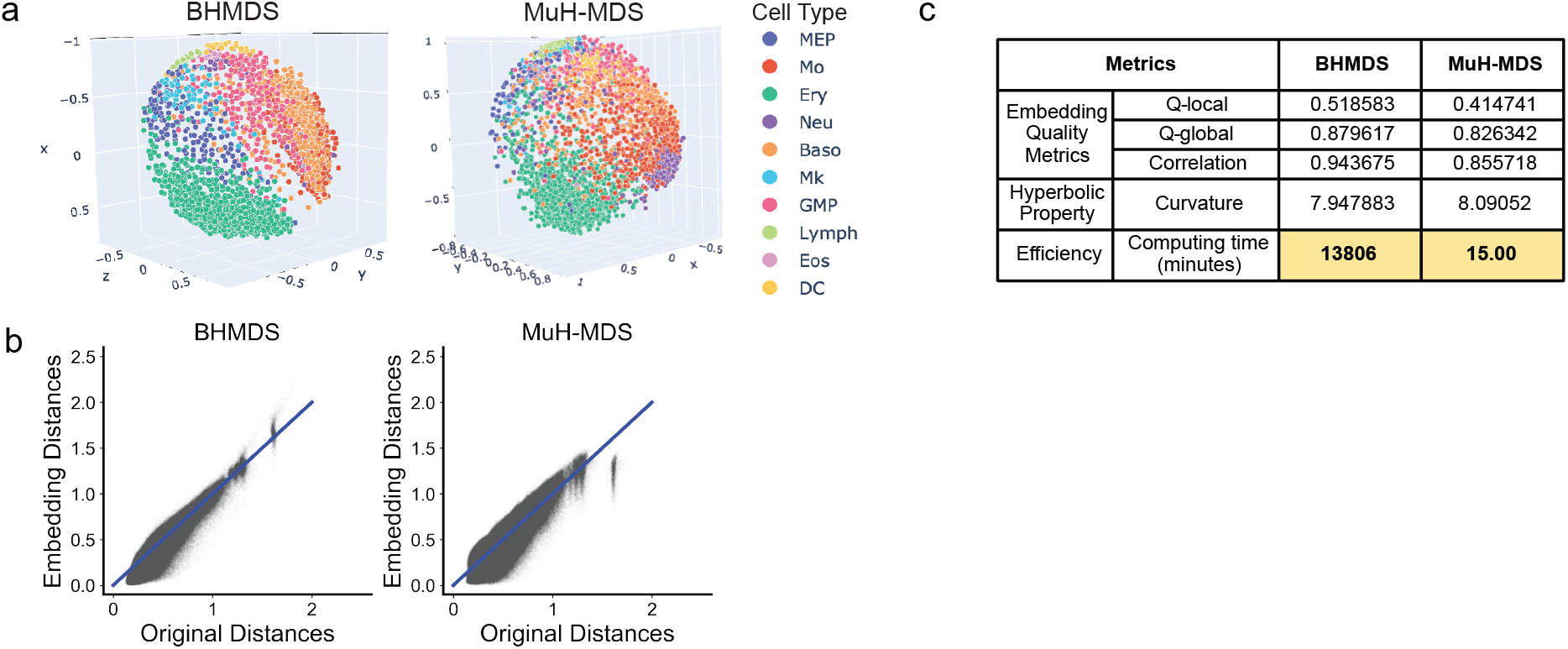
Comparison between the original (BHMDS) and multiscale (MuH-MDS) approaches using the mouse myeloid progenitors scRNA-seq dataset (“Paul”) [2]. (a) Shepard diagram comparing original pairwise distances with embedded distances. (b) Embedding in 3-d hyperbolic space of “Paul” dataset. (c) Embedding metrics. MuH-MDS parameters: *α* = 30, *k* = 210.

**Figure S2:**
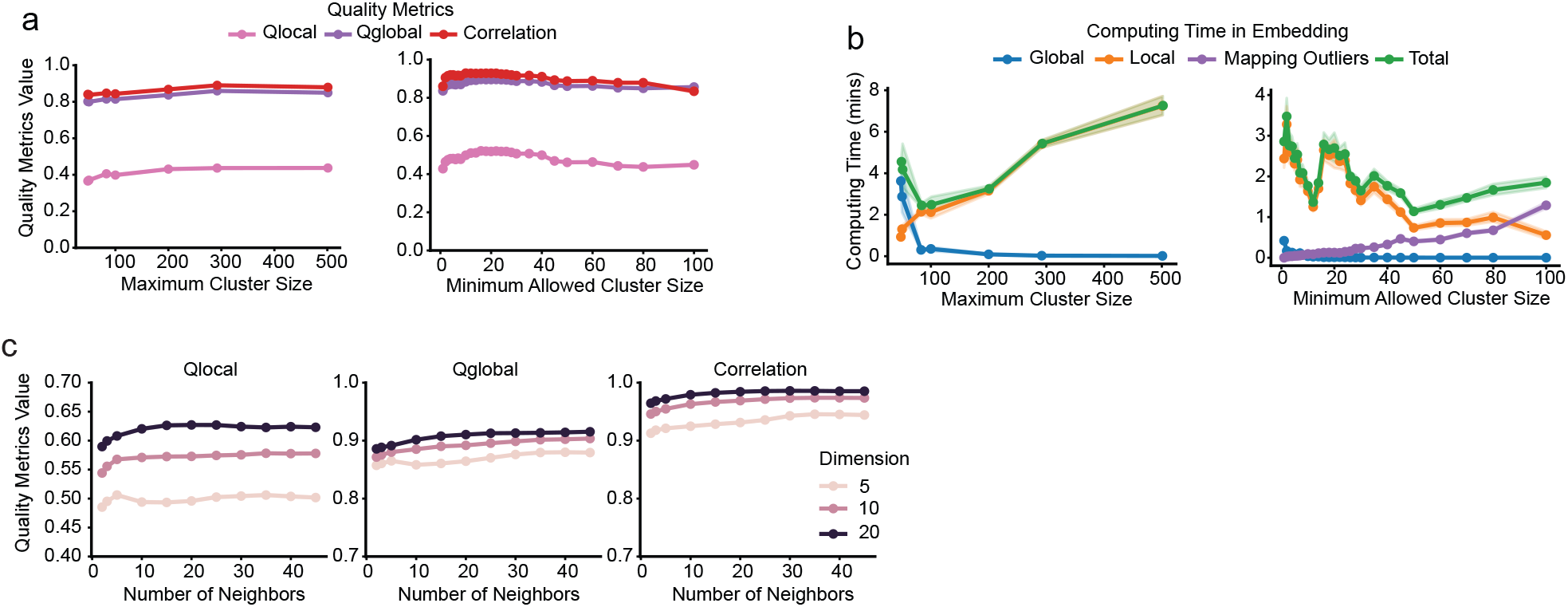
Simulation of hyperparameters using the Paul dataset [2], with 20 repetitions for each parameter setting. (a-b) Changes in embedding quality metrics (a) and running time (b) with varying maximum (left) or minimum (right) population sizes. (c) Embedding quality metrics across different numbers of nearest neighbors (*α*) in dimensions 5, 10, and 20. MuH-MDS parameters: (1) For maximum cluster size simulations: *k* = 25 *α* = 20 *D* = 3; (2) For minimum allowed cluster size simulations: *k* = 100 *α* = 20 *D* = 3; (3) For *α* and *D* simulations: *k* = 50.

**Figure S3:**
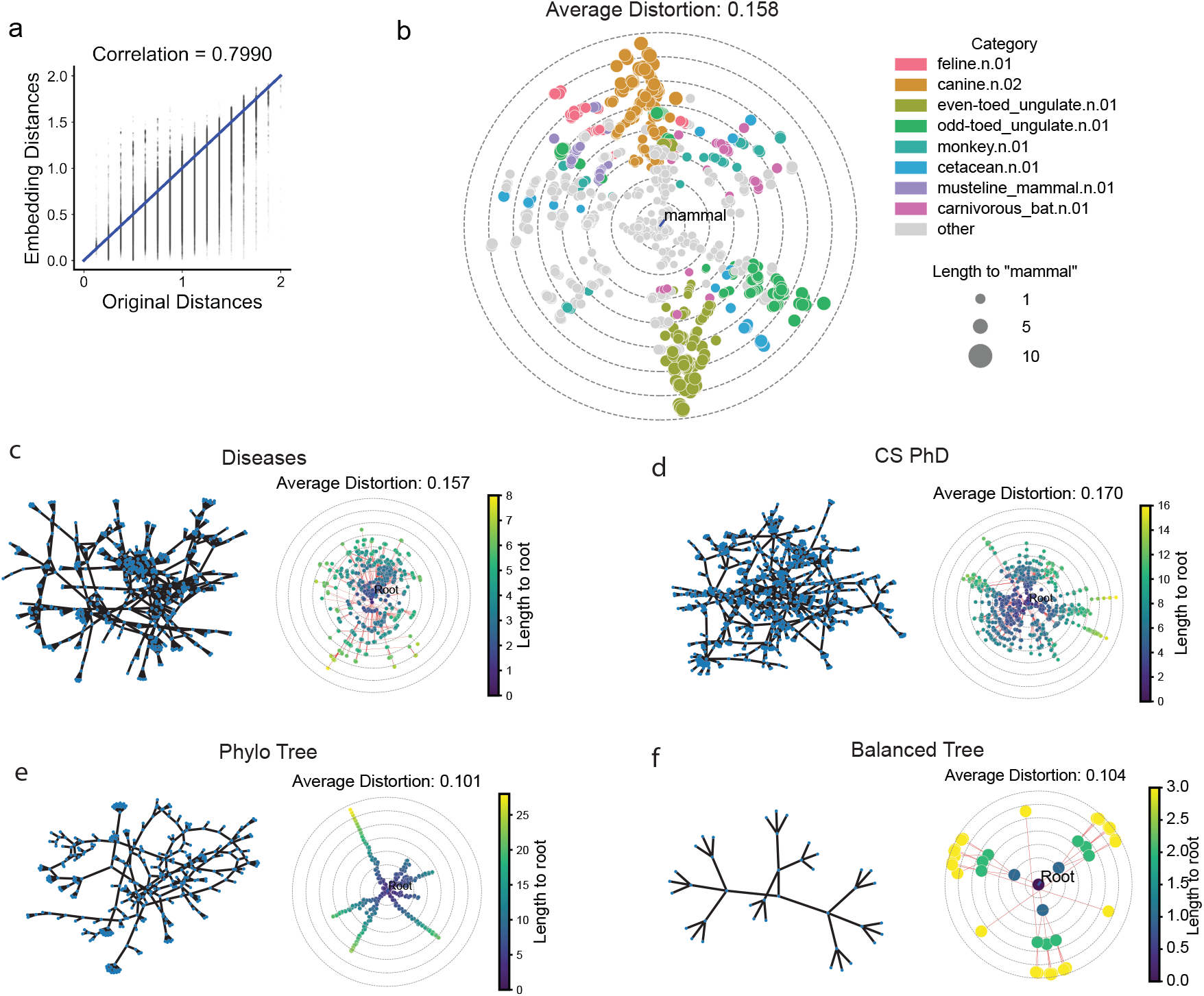
(a) Shepard diagram of WordNet mammal subtree embedding in 2-d hyperbolic space. (b) Application of MuH-MDS to the WordNet mammal subtree, shown in native coordinates. Marker size reflects the graph distance to the root node “mammal”, while color denotes membership in semantic categories as shown in the legend. These categories correspond to parent nodes with the largest numbers of child nodes and were selected to highlight major structural branches within the hierarchy. MuH-MDS parameters: *k* = 198, *α* = 60. (c-f) Embedding of multiple graph datasets from De Sa et al.[3] using MuH-MDS into 2D hyperbolic space. Left: original graph representations. Right: embedding results in 2D hyperbolic space. Color indicates graph distance to the root. Root is defined as the node with the smallest average distance to all other nodes in the original graph. MuH-MDS parameters: Diseases: *k* = 106, *α* = 60, CS PhD: *k* = 75, *α* = 60, Phylo Tree: *k* = 34, *α* = 20, Balanced Tree: *k* = 40, *α* = 20. For comparison, the distortion rates obtained by HoroPCA are: Diseases: 0.15, CS PhD: 0.16, Phylo Tree: 0.13, Balanced Tree: 0.19. MuH-MDS obtained comparable or better results on each dataset.

**Figure S4:**
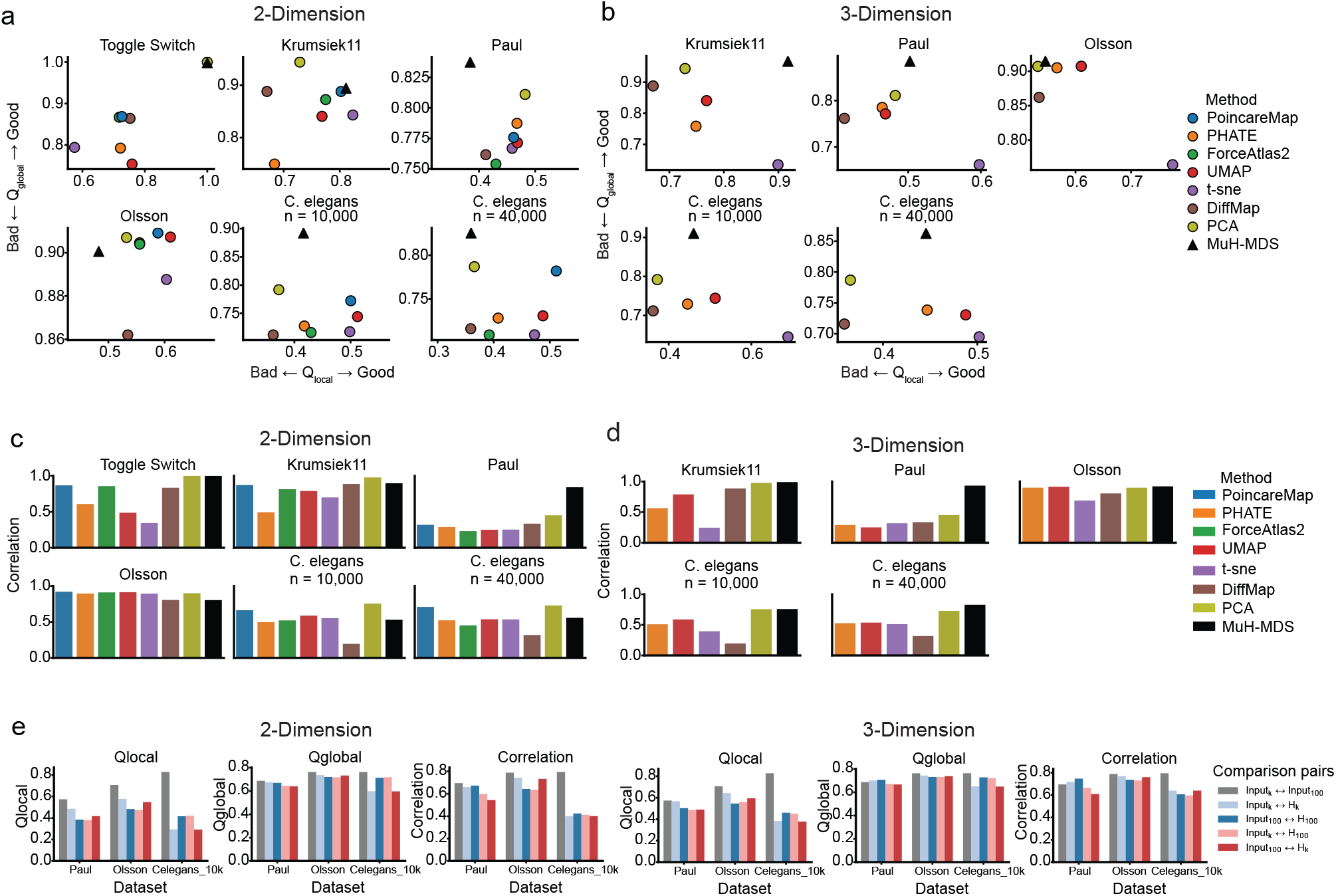
(a-d) Embedding quality metric for all datasets tested in the paper, in 2-dimension (a-b) or 3-dimension (c-d) space. MuH-MDS parameters: Toggle switch, 2d: *α* = 10,*k* = 20. Krumsiek11, 2d:*α* = 10,*k* = 50; 3d: *α* = 20,*k* = 50. Paul, 2d and 3d: *α* = 20,*k* = 100. Olsson, 2d: *α* = 30,*k* = 40; 3d: *α* = 20,*k* = 40. *C. elegans* (10,000), 2d: *α* = 20,*k* = 300; 3d: *α* = 30,*k* = 300. *C. elegans* (40,000), 2d and 3d: *α* = 30,*k* = 200. (*α*: number of neighbors, *k*: number of clusters) (e) Robustness of representation structure to PCA dimensionality in preprocessing. Bars show pairwise embedding quality scores computed between different representation pairs. Input_*k*_ and Input_100_ denote PCA-reduced Euclidean input spaces retaining *k* and 100 principal components, respectively (*k* = 20 for Paul and Olsson, and *k* = 60 for *C. elegans*). H_*k*_ and H_100_ denote hyperbolic embeddings learned from Input_*k*_ and Input_100_. Matched (Input_*k*_ H_*k*_, ↔ Input_100_ ↔ H_100_) and cross (Input_*k*_ ↔ H_100_, Input_100_ ↔ H_*k*_) comparisons assess the sensitivity of the learned representations to the number of retained PCs. Similar values observed for matched and cross comparisons indicate that the learned hyperbolic representations are robust to the choice of PCA dimensionality.

**Figure S5:**
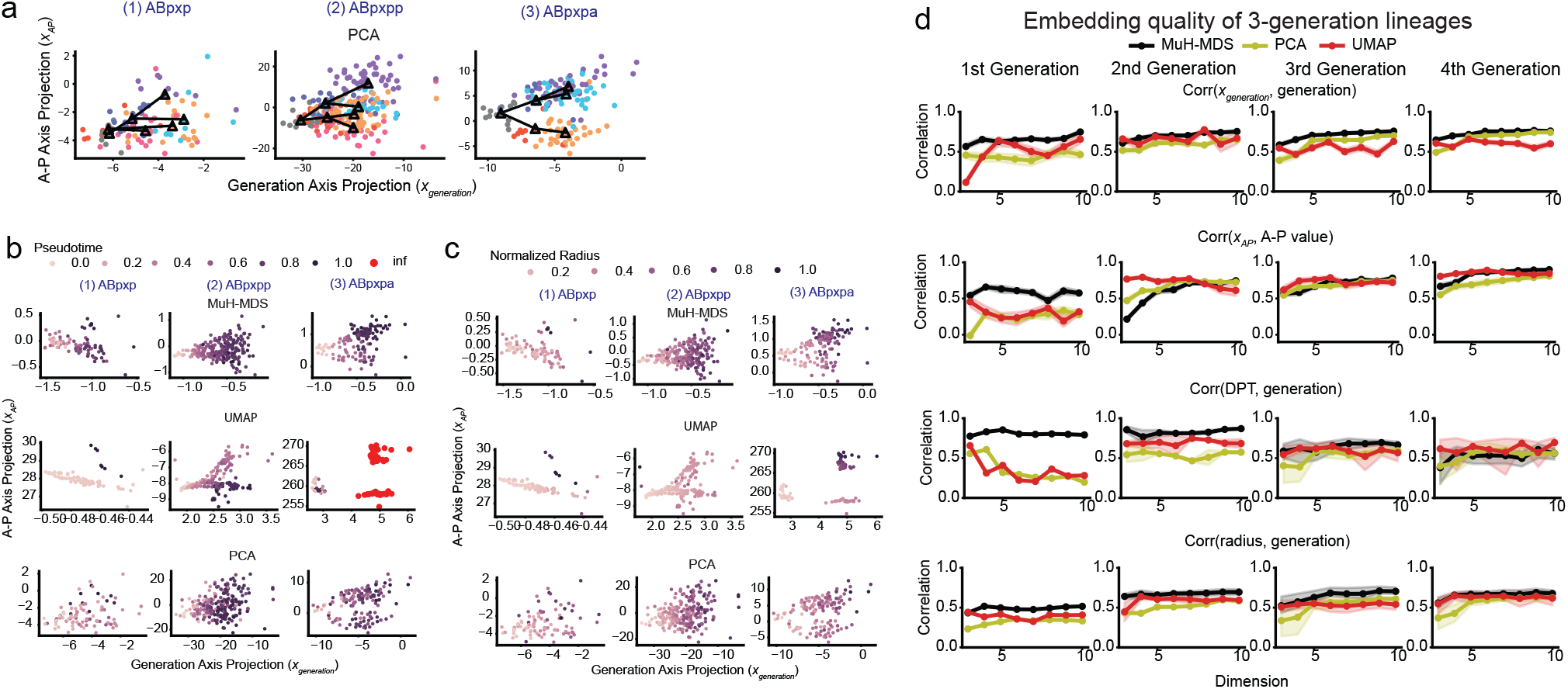
Analysis of the ABpxp lineage (7,562 samples) of *C. elegans* dataset. (a) 7-d PCA projecting to 2-d space, for 3-generation lineages starting from ABpxp, ABpxpp and ABpxpa. (b) Pseudotime computed from 7-d embedding of MuH-MDS, UMAP and PCA, for 3-generation lineages starting from ABpxp, ABpxpp and ABpxpa respectively. Some samples have infinite pseudotime because graph could be disconnected in high dimensional UMAP. (c) Normalize, re-centered radius computed from same embeddings as in (b). All embeddings are re-centered so that its population centroid is at the origin before computing the radius. (d) Comparison of 4 sub-lineage embedding quality metrics: (1) correlation between generation and projected coordinates (Corr(*x*_generation_, generation)), (2) correlation between A-P value and projected coordinates (Corr(*x*_AP_, A-P value)), (3) correlation between generation and diffusion pseudotime inferred from embedding (Corr(DPT, generation)) and (4) correlation between generation and sample radius in embeddings (Corr(radius, generation)), across different dimensions, embedding methods (MuH-MDS, PCA, and UMAP), for lineages starting from 1st generation to 4th generation.

**Figure S6:**
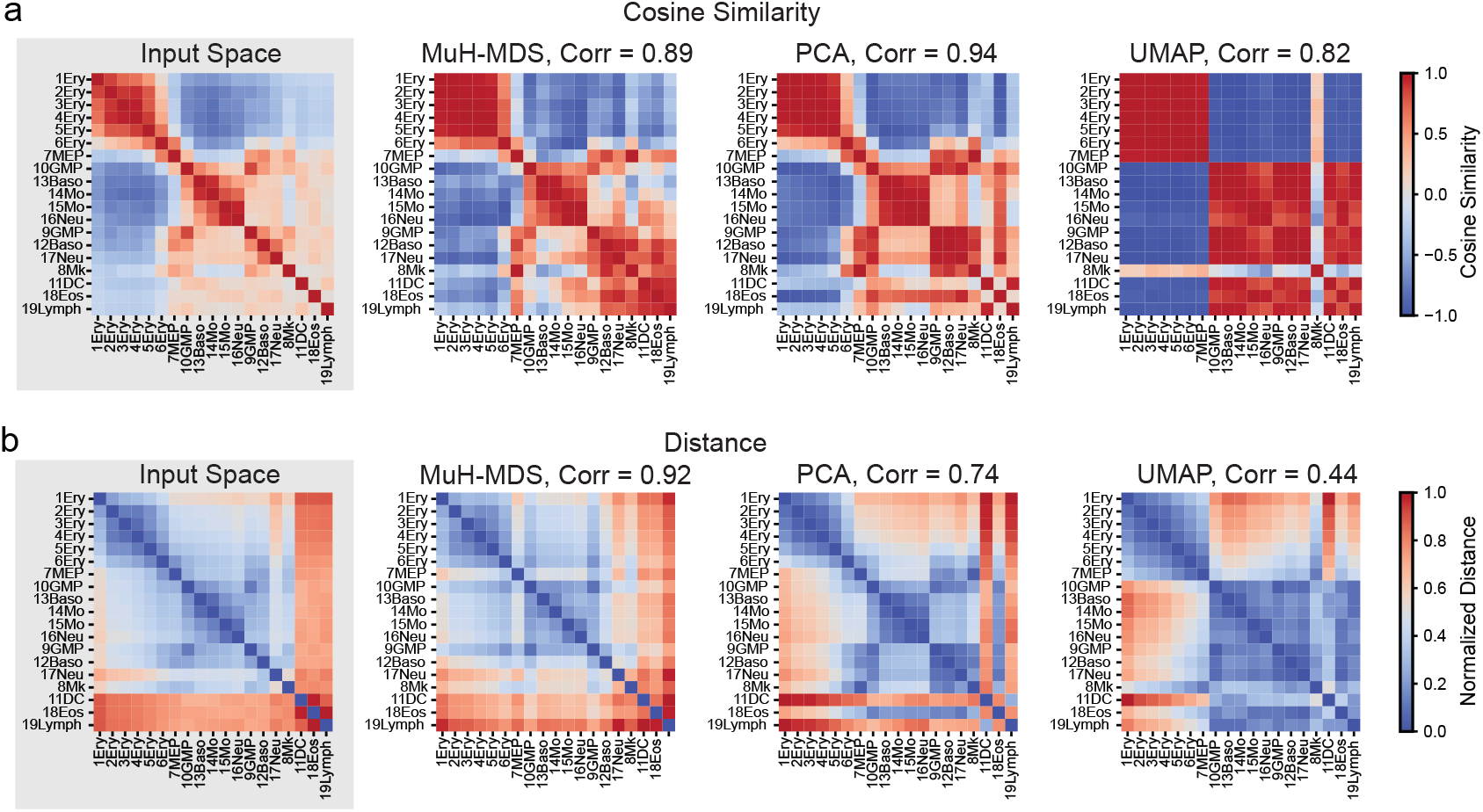
(a-b) Geometric comparison between the input space (100-PC PCA space derived from the original RNA-seq data) and each three-dimensional embedding, for all 19 clusters in “Paul” dataset. MuH-MDS best preserves both (a) cosine similarity and (b) pairwise distances from the input space.

**Figure S7:**
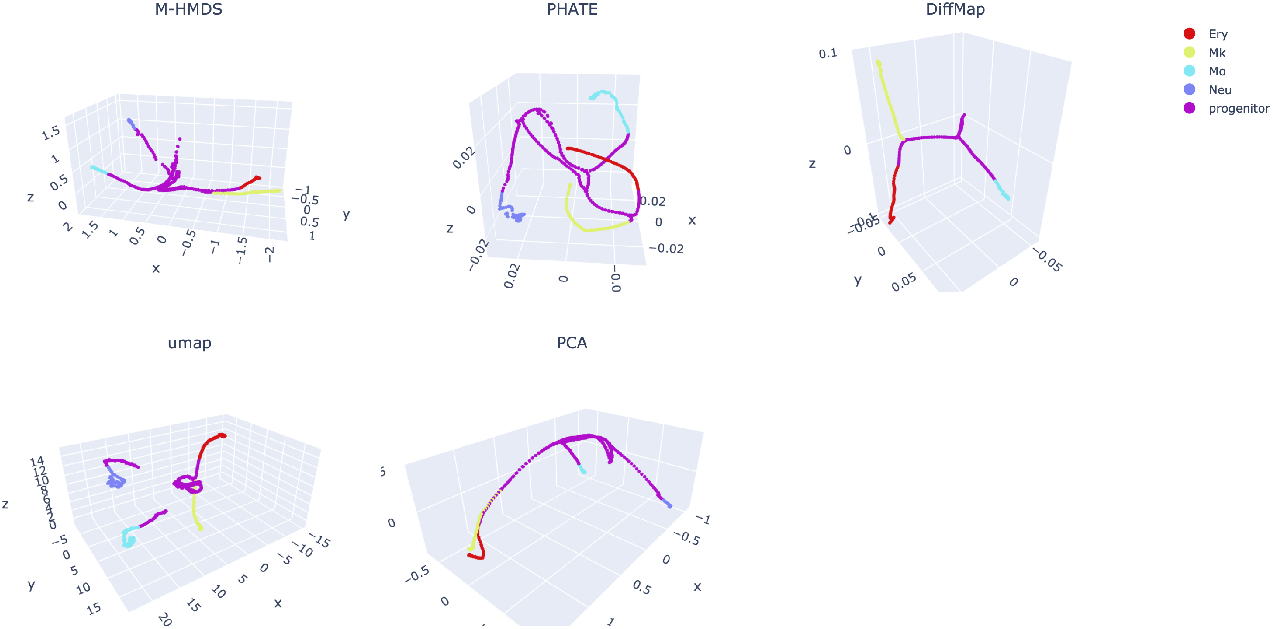
Embedding of the (synthetic) myeloid progenitors dataset (Krumsiek11), in 3-d space.

**Figure S8:**
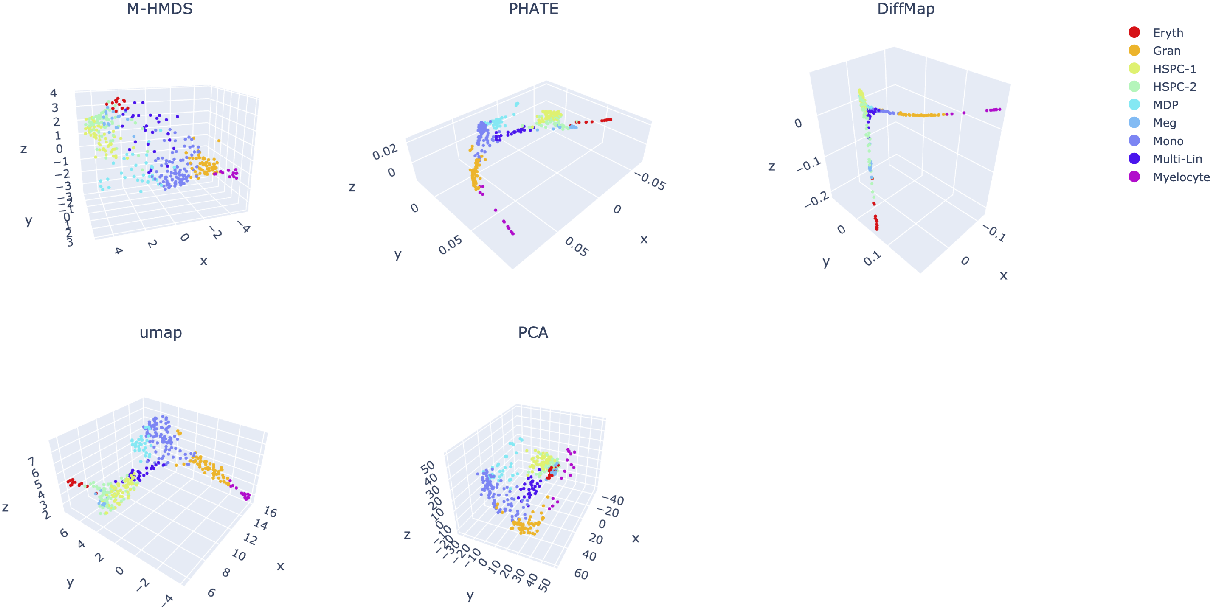
Embedding of the mouse myelopoesis dataset (Olsson), in 3-d space.

**Figure S9:**
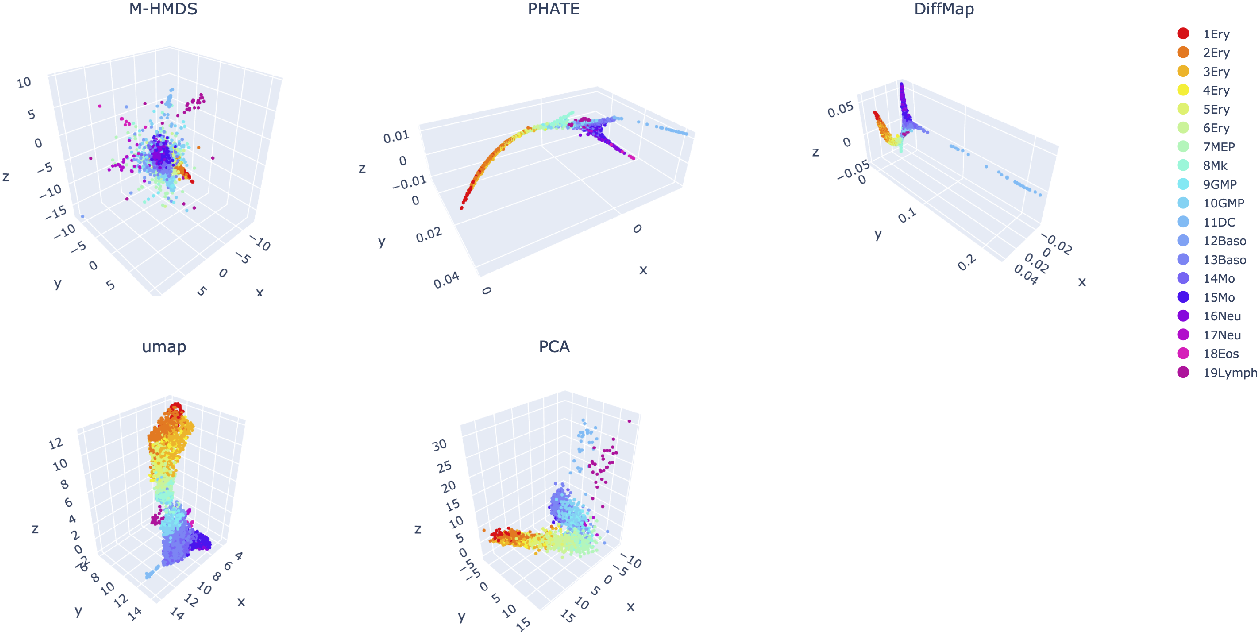
Embedding of the mouse myeloid progenitors dataset (Paul), in 3-d space.

**Figure S10:**
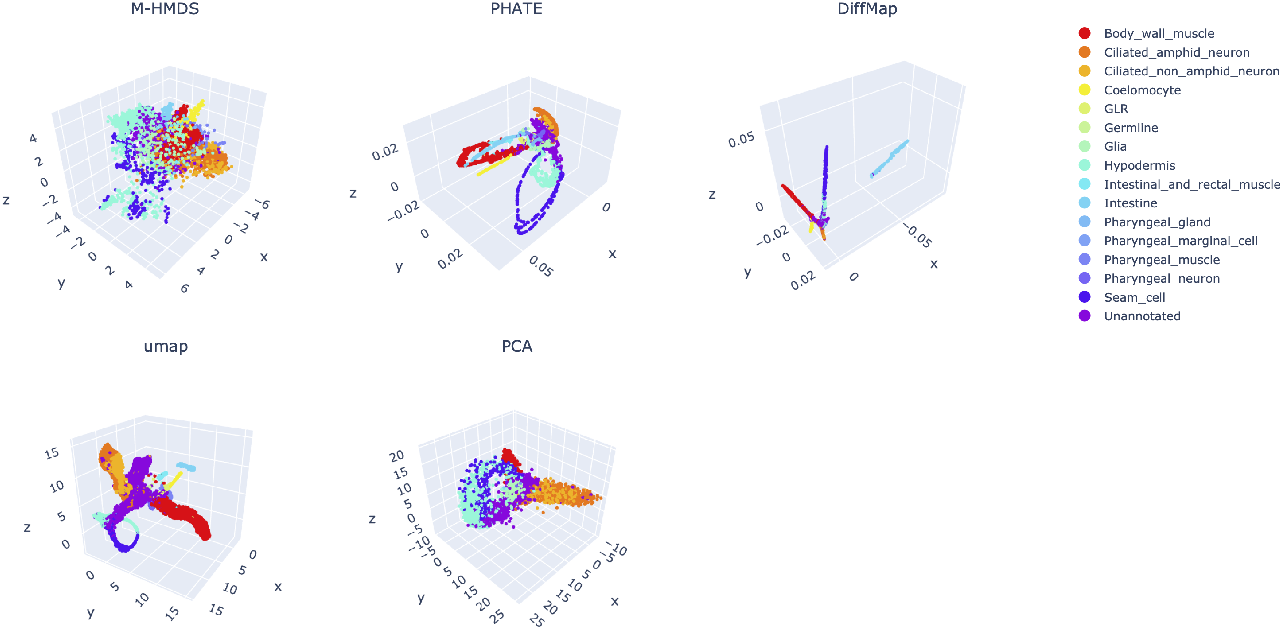
Embedding of the 10,000 sample subset of *C. elegans* dataset, in 3-d space, colored by cell types.

**Figure S11:**
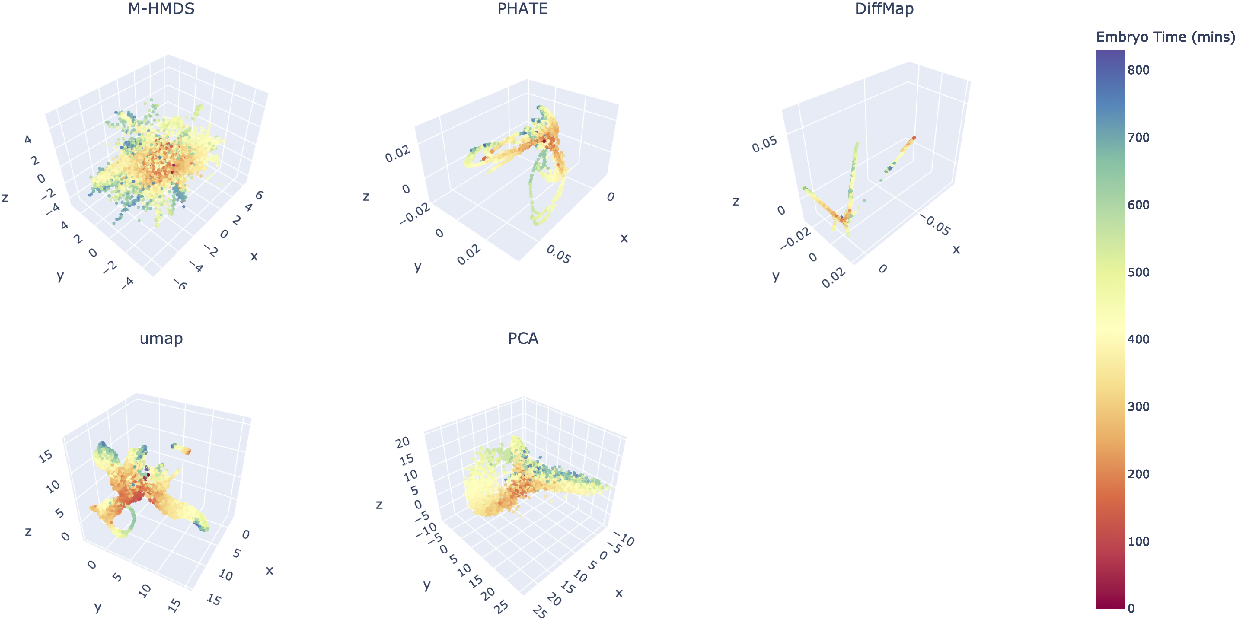
Same as Fig. (S10), but colored by embryo time.

**Figure S12:**
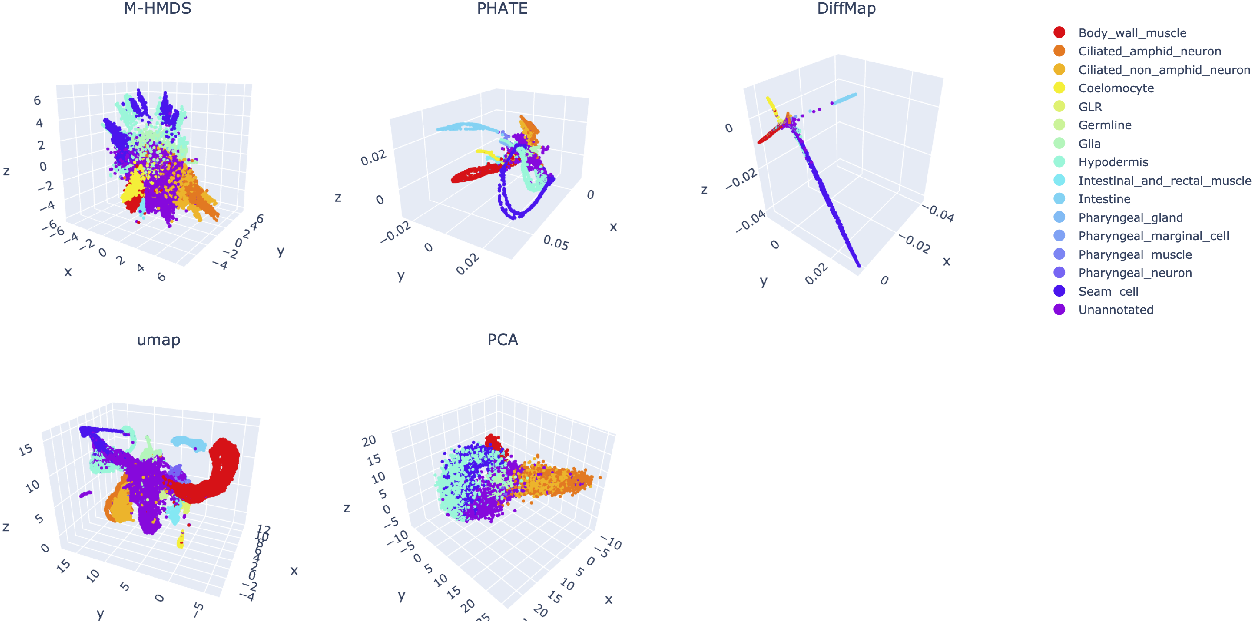
Embedding of the 40,000 sample subset of *C. elegans* dataset, in 3-d space, colored by cell types.

**Figure S13:**
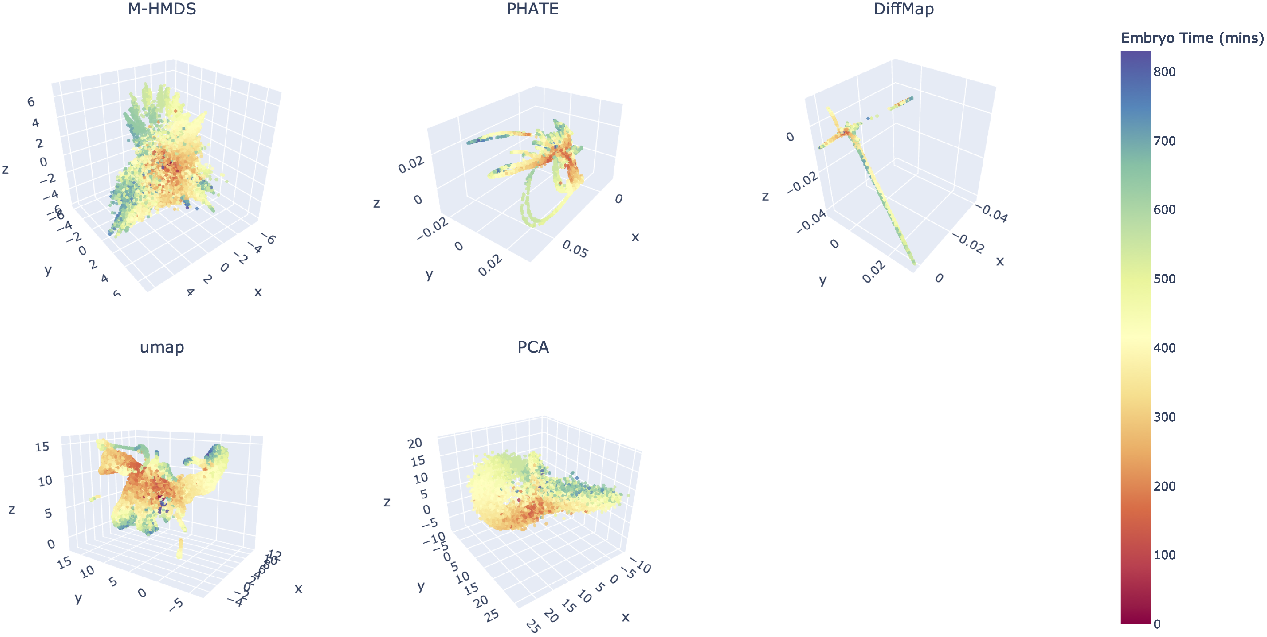
Same as Fig. (S12), but colored by embryo time.

**Figure S14:**
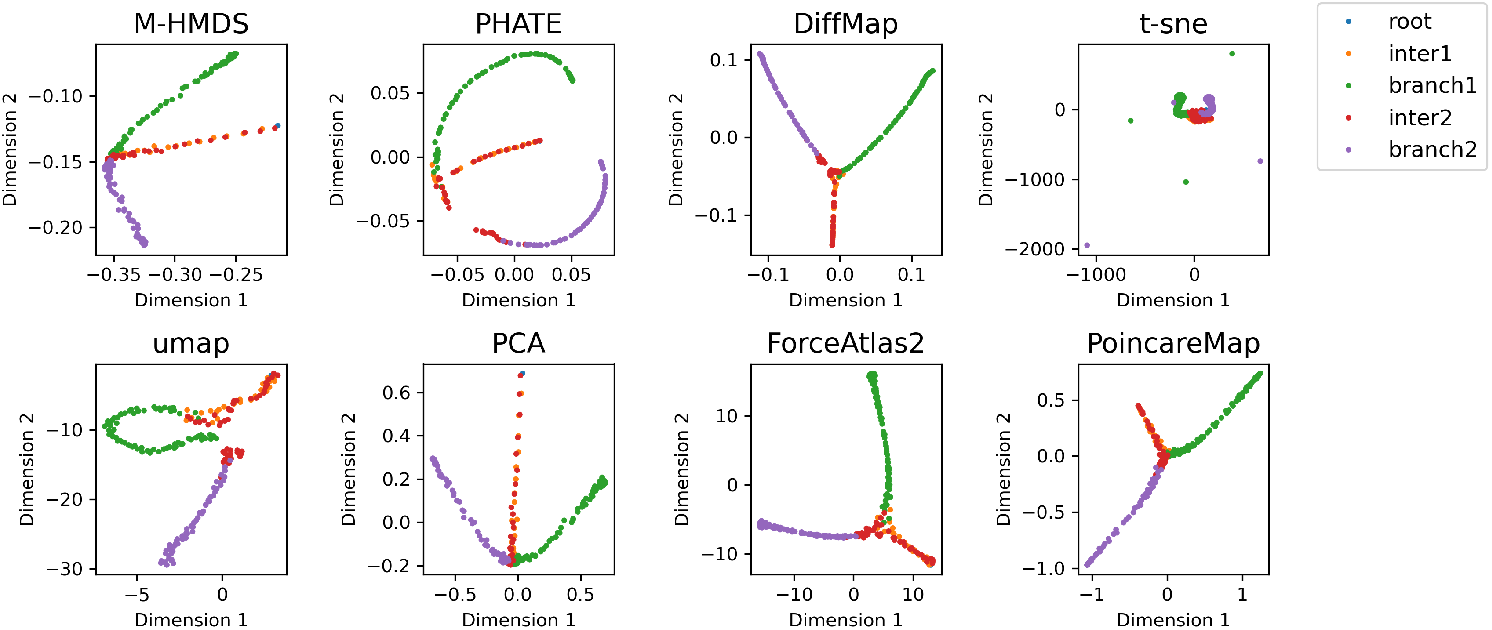
Embedding of the (synthetic) toggle switch dataset, in 2-d space.

**Figure S15:**
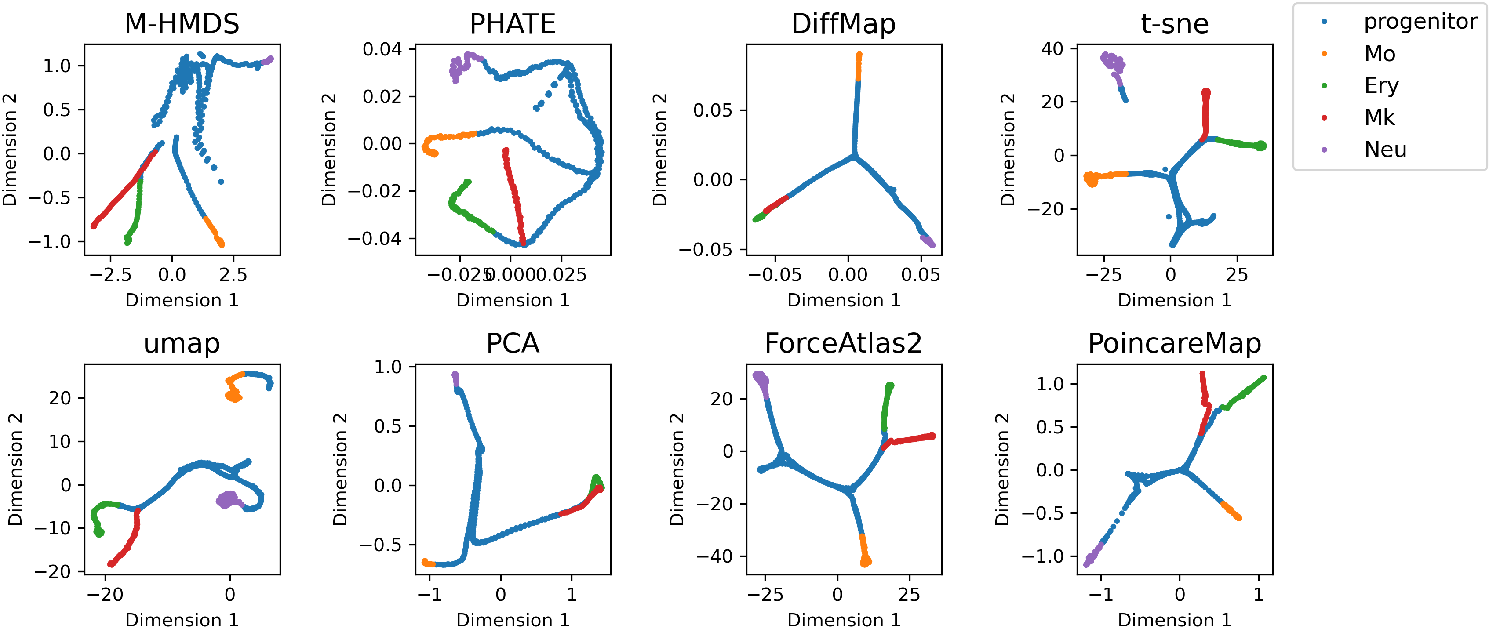
Embedding of the (synthetic) myeloid progenitors dataset (Krumsiek11), in 2-d space.

**Figure S16:**
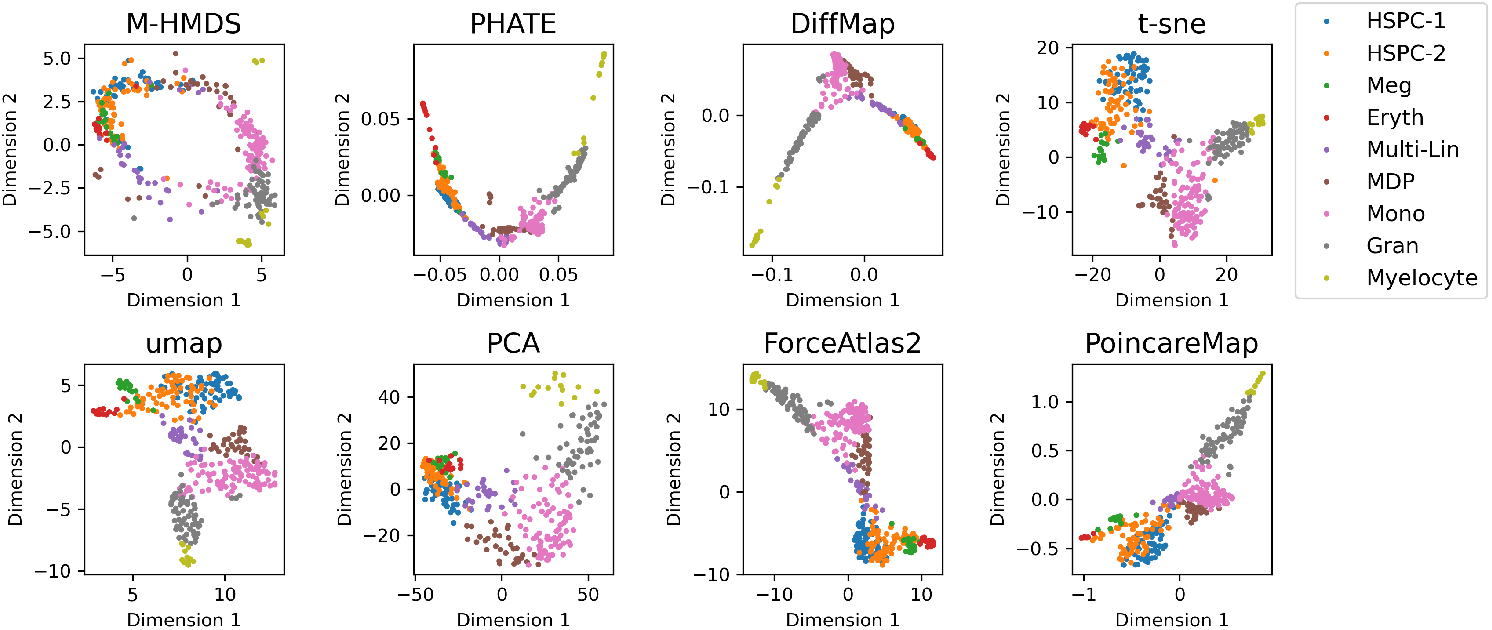
Embedding of the mouse myelopoesis dataset (Olsson), in 2-d space.

**Figure S17:**
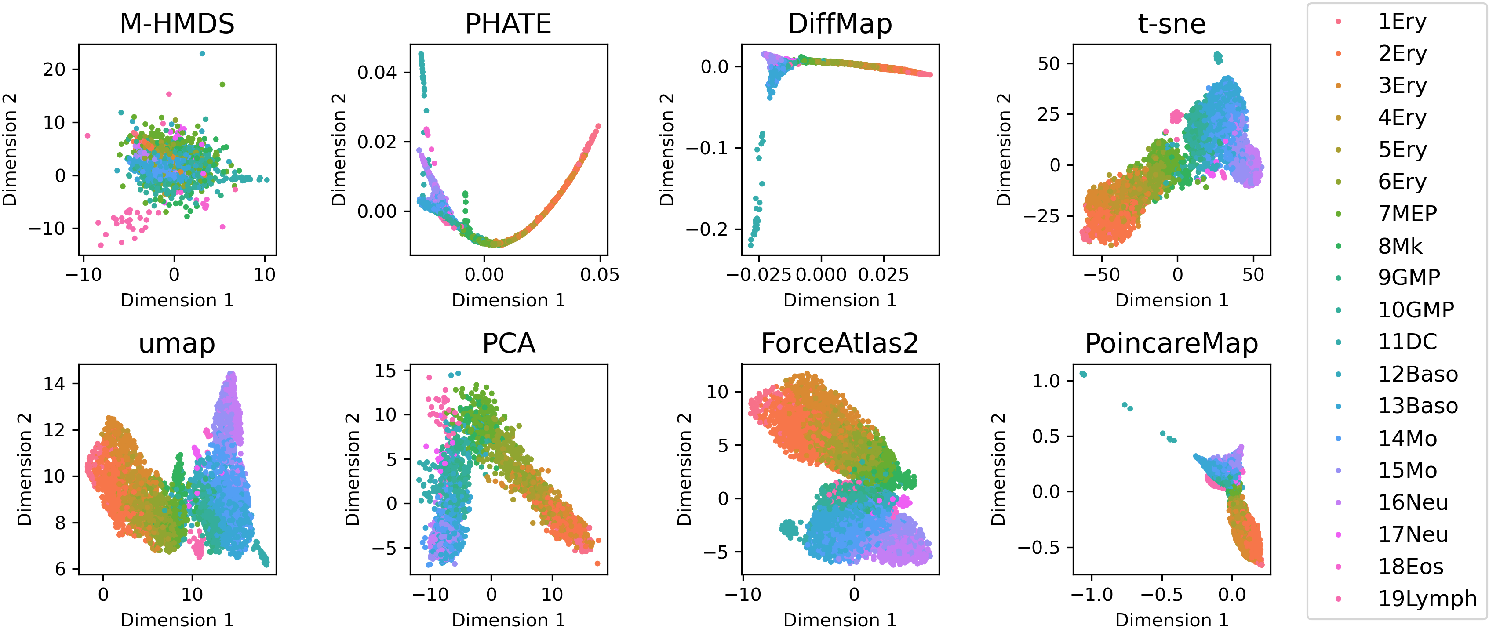
Embedding of the mouse myeloid progenitors dataset (Paul), in 2-d space.

**Figure S18:**
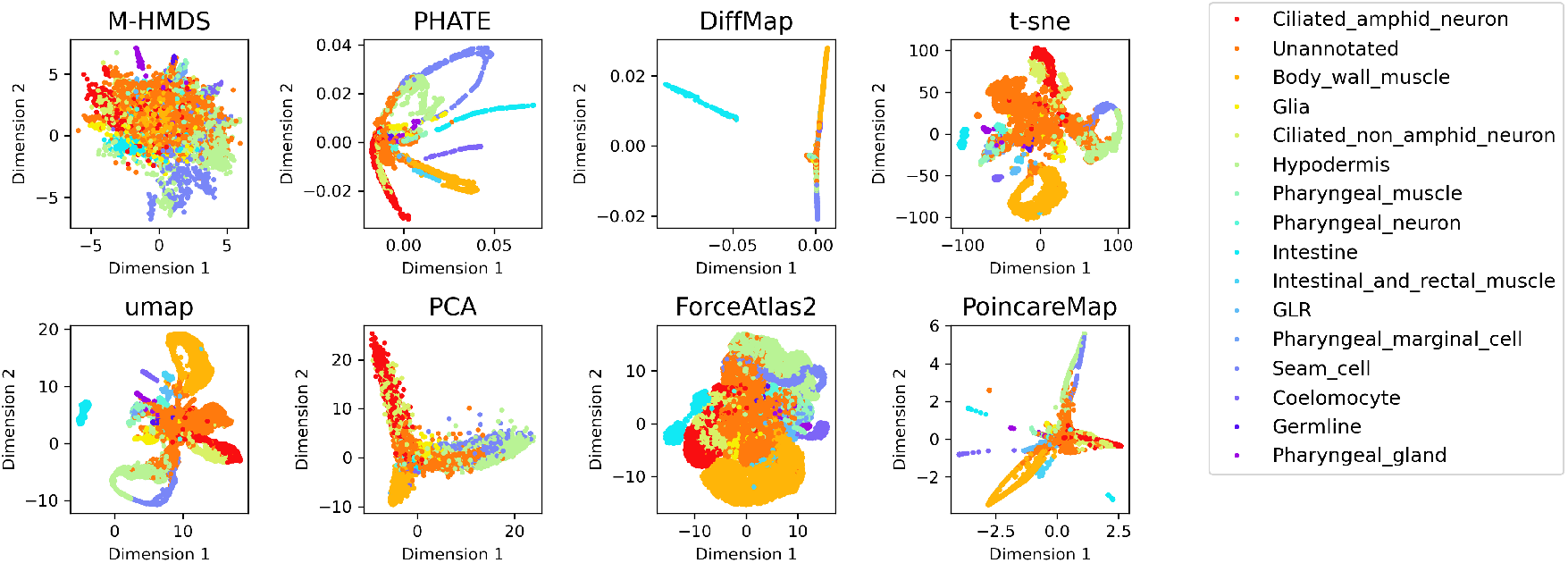
Embedding of the 10,000 sample subset of *C. elegans* dataset, in 2-d space, colored by cell types.

**Figure S19:**
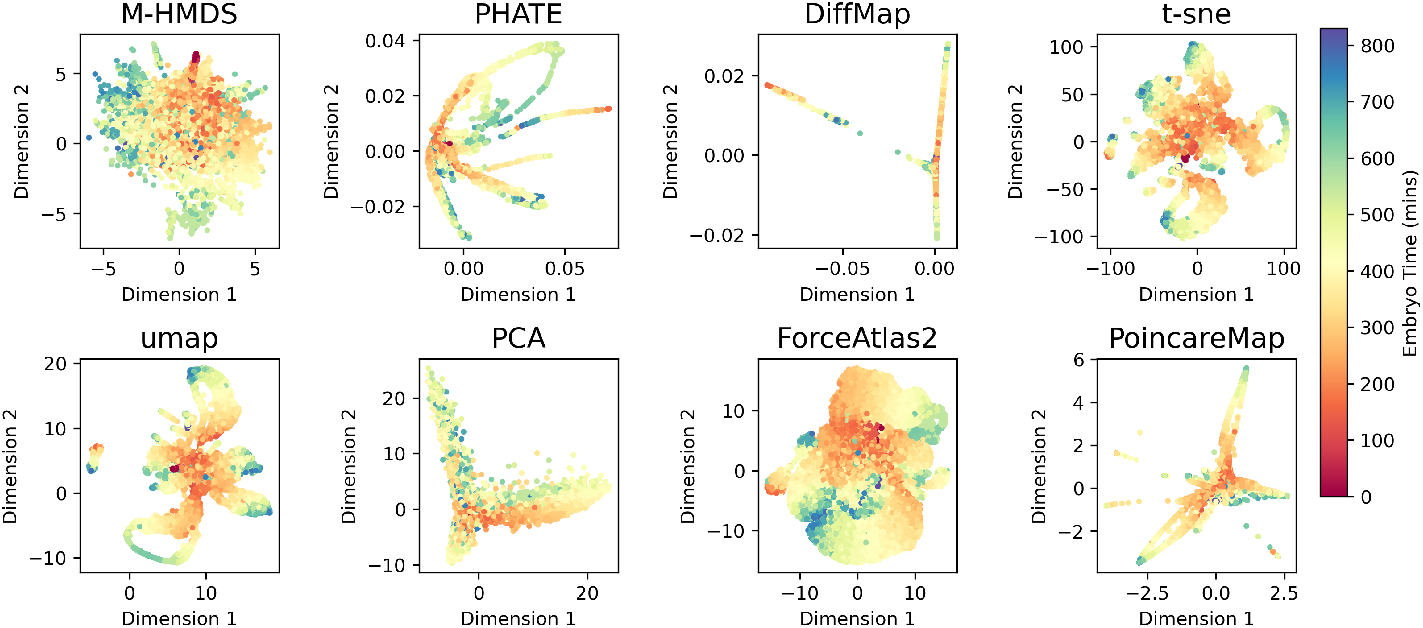
Same as Fig. (S18), but colored by embryo time.

**Figure S20:**
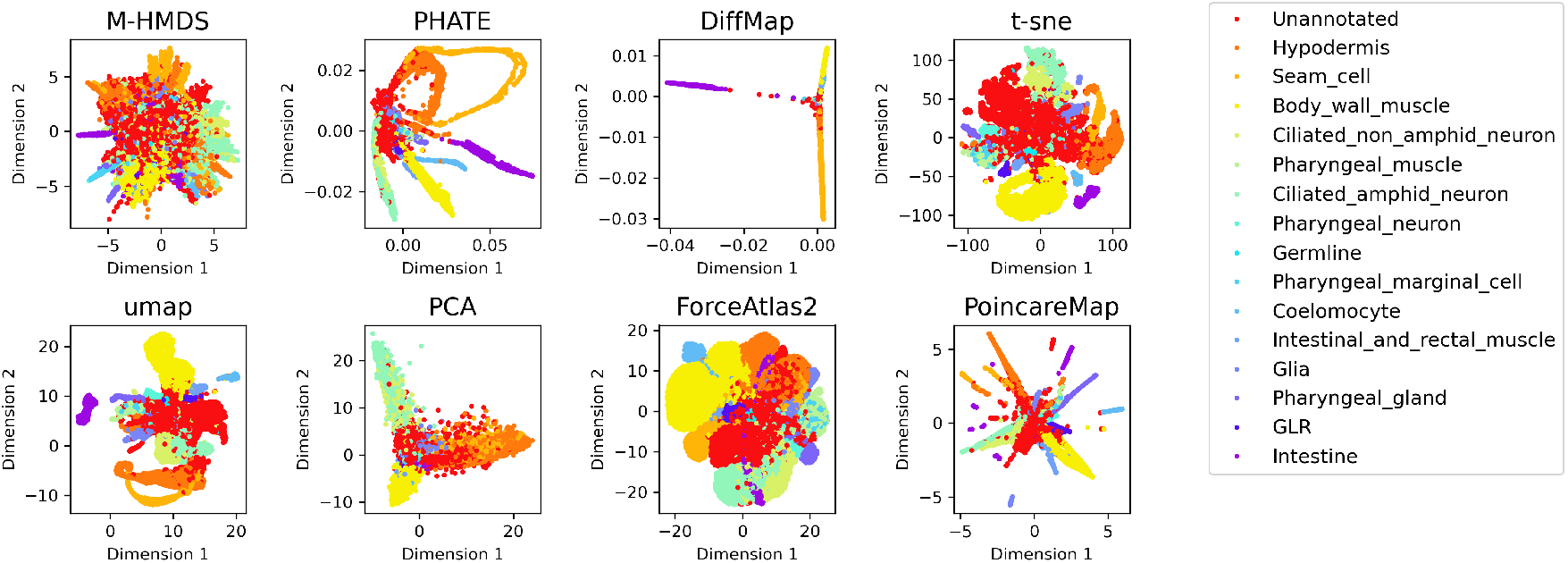
Embedding of the 40,000 sample subset of *C. elegans* dataset, in 2-d space, colored by cell types.

**Figure S21:**
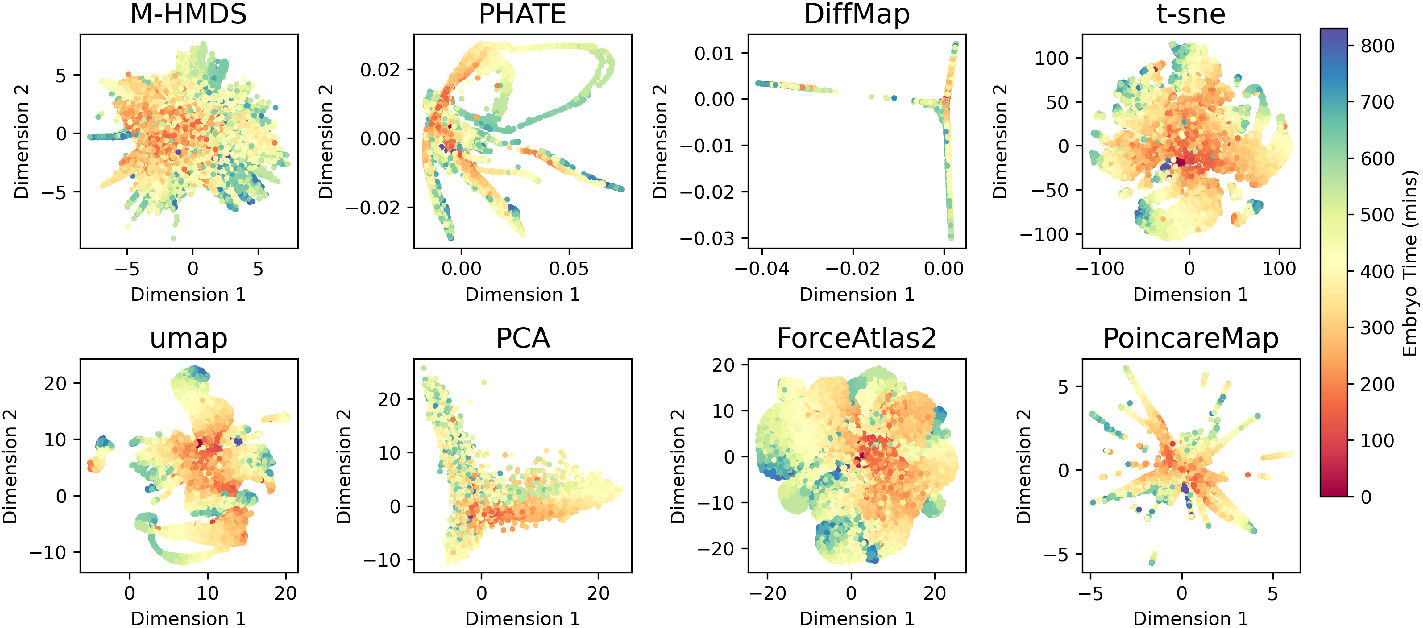
Same as Fig. (S20), but colored by embryo time.

## Notes

### Competing Interest Statement

The authors have declared no competing interest.

### Summary of Updates

Updated manuscript with additional figure: figure 5. Some modifications to wording of title, abstract and introduction.

